# Site-specific ubiquitination of the E3 ligase HOIP regulates cell death and immune signaling

**DOI:** 10.1101/742544

**Authors:** Lilian M. Fennell, Luiza Deszcz, Alexander Schleiffer, Karl Mechtler, Anoop Kavirayani, Fumiyo Ikeda

**Author notes:** Correspondence: Fumiyo Ikeda.

## Abstract

HOIP, the catalytic component of the Linear Ubiquitin chain Assembly Complex (LUBAC), is a critical regulator of inflammation. However, how HOIP itself is regulated to control inflammatory responses is unclear. Here, we discover that site-specific ubiquitination of K784 within HOIP promotes Tumour Necrosis Factor (TNF)-induced inflammatory signalling by controlling TNF Receptor complex I (TNFR1) formation. A HOIP K784R mutant is catalytically active but shows reduced induction of an NF-κB reporter relative to wild type HOIP. HOIP K784 is evolutionarily conserved, equivalent to HOIP K778 in mice. We generated *Hoip^K778R/K778R^* knockin mice, which show no overt developmental phenotypes; however, in response to TNF, *Hoip^K778R/K778R^* mouse embryonic fibroblasts display suppressed NF-κB activation and increased sensitivity to apoptosis. On the other hand, HOIP K778R enhances the TNF-induced formation of TNFR complex II, and an interaction between TNFR complex II and LUBAC. Loss of the LUBAC component SHARPIN leads to embryonic lethality in *Hoip^K778R/K778R^* mice, which is rescued by knockout of TNFR1. We propose that site-specific ubiquitination of HOIP regulates a LUBAC-dependent switch between survival and apoptosis in TNF-signalling.

## Introduction

The Linear UBiquitin chain Assembly Complex (LUBAC) is a critical regulator of inflammation in humans and mice (Ikeda, 2015, Peltzer & Walczak, 2019, Sasaki & Iwai, 2015, Walczak, 2011). LUBAC influences the inflammatory response by regulating the tumour necrosis factor (TNF)-signaling pathway. Upon TNF binding, the TNF receptor (TNFR) forms TNFR complex I (TNFR1), consisting of TNF Receptor type 1-Associated DEATH Domain (TRADD), Receptor-Interacting serine/threonine-Protein Kinase 1 (RIPK1), TNF Receptor-Associated Factor 2 (TRAF2), Cellular Inhibitor of Apoptosis Protein (cIAP) 1/2 and LUBAC. TNFR1 complex I promotes cell survival via downstream signaling cascades such as NF-κB through the key kinase complex IκB kinase (IKK) consisting of IKK1/2 and NF-κB Essential Modifier (NEMO). Post-translational modifications, including ubiquitination, regulate multiple events in this signaling cascade. Linear/Met1-, Lys11- and Lys63-ubiquitin linkage types regulate the recruitment of specific signaling complexes (Peltzer et al., 2016, Witt & Vucic, 2017), whereas Lys48-linked ubiquitin chains trigger degradation by the ubiquitin-proteasome system. As part of TNFR complex I, LUBAC generates linear/Met1-ubiquitin chains on NEMO (Tokunaga et al, Haas et al 2009) and RIPK1 (Gerlach et al., 2011) to promote NF-κB signaling. NF-κB activation leads to gene inductions of anti-apoptosis genes such as Cellular FLICE-like inhibitory Protein (cFLIP), thus known as anti-apoptosis pathway (Lamkanfi et al., 2007, Peltzer & Walczak, 2019).

On the other hand, when NF-κB pathway is disturbed, TNF can promote apoptosis via formation of the TNFR complex II, which consists of RIPK1, TRADD, FAS-Associated Death Domain (FADD) and Caspase 8 (Justus & Ting, 2015, Witt & Vucic, 2017). TNFR complex II formation also appears to be regulated by LUBAC (Asaoka & Ikeda, 2015, Peltzer & Walczak, 2019, Sasaki & Iwai, 2015), but the mechanisms are unclear.

LUBAC consists of the E3 ligase HOIP/RNF31, and two subunits HOIL-1L/RBCK1 and SHARPIN/SIPL1 (Gerlach et al., 2011, Ikeda et al., 2011, Rittinger & Ikeda, 2017, Tokunaga et al., 2011). Genetic loss of HOIP or HOIL-1L triggers embryonic lethality in mice due to upregulation of apoptosis, uncovering their essential roles in mouse embryonic development and cell death regulation (Emmerich et al., 2013, Hrdinka & Gyrd-Hansen, 2017, Peltzer et al., 2014) (Meier et al., 2015). In contrast, SHARPIN-deficient mice (*Sharpin^cpdm/cpdm^*) suffer from systemic inflammation accompanied with chronic proliferative dermatitis at the age of 6-8 weeks (Kumari et al., 2014, Seymour et al., 2007). Skin tissues derived from *Sharpin^cpdm/cpdm^* mice show immune cell infiltrations and upregulation of keratinocyte apoptosis (Seymour et al., 2007). The phenotypes of these genetically modified mice are at least partially rescued by TNFR1 knockout, suggesting that LUBAC attenuates apoptosis downstream of the TNF signaling cascade (Kumari et al., 2014, Rickard et al., 2014). HOIL-1L and HOIP mutations are observed in patients with autoimmune diseases, implicating LUBAC in the regulation of immune responses in humans (Boisson et al., 2015, Boisson et al., 2012).

At the molecular level, HOIP is a RING-IBR-RING (RBR) type of E3 ligase that specifically generates linear/Met1-linked ubiquitin chains with SHARPIN and HOIL-1L. Linear ubiquitin chains are atypical chains linked via the C-terminal Gly of one ubiquitin moiety to the N-terminal Met1 of another ubiquitin moiety. The catalytic center of HOIP is in the second RING domain (Stieglitz et al., 2012), and the Linear Ub chain Determining Domain (LDD) domain provides its unique ability to generate linear ubiquitin chains (Smit et al., 2012). Thus far, HOIP is the only ligase known to generate linear ubiquitin chains (Dove & Klevit, 2017, Kirisako et al., 2006).

*In vitro*, HOIP requires HOIL-1L or SHARPIN to generate linear ubiquitin chains (Gerlach et al., 2011, Ikeda et al., 2011, Kirisako et al., 2006, Tokunaga et al., 2011). However, the HOIP RBR-LDD fragment is active in the absence of HOIL-1L and SHARPIN, suggesting a self-inhibitory mechanism (Smit & Sixma, 2014, Walden & Rittinger, 2018). LUBAC generates linear/Met1 ubiquitin chains at Lys on substrates, which depends on HOIL-1L (Smit et al., 2013). In cells, HOIP exists mostly in complex with SHARPIN or HOIL-1L (Kirisako et al., 2006, Tokunaga et al., 2011), thus, the LUBAC complex is expected to be active. Yet, the LUBAC-dependent downstream cascades are dependent on stimuli like TNF. The mechanisms that regulate LUBAC activity are unclear. In particular, it is not known how inflammatory stimuli modulate the interactions between LUBAC and its substrates.

Two deubiquitinases (DUBs), called “OTU DUB with LINear linkage specificity” (OTULIN) and CYLD, hydrolyze linear ubiquitin chains and regulate inflammatory signaling cascades. Both OTULIN and CYLD can form a complex with HOIP (Elliott et al., 2016, Elliott et al., 2014, Fiil et al., 2013, Hrdinka et al., 2016, Keusekotten et al., 2013, Kupka et al., 2016, Schaeffer et al., 2014, Takiuchi et al., 2014, Wagner et al., 2016). However, loss-of-function of OTULIN and CYLD in mice does not resulted in expected phenotypes compared with LUBAC-deficient mice (Damgaard et al., 2016, Reiley et al., 2006, Zhang et al., 2006); knockin mice expressing OTULIN C129A (a dominant negative mutant) are embryonic lethal with increased apoptosis signals, partially overlapping with mouse phenotype of HOIP and HOIL-1L knockout (Heger et al., 2018, Peltzer et al., 2018, Peltzer et al., 2014). Recently, it was shown that the OTULIN mutant Cys129Ala increases ubiquitination signal of all three LUBAC components (Heger et al., 2018). Hyper-ubiquitinated LUBAC in the OTULIN mutant Cys129Ala expressing cells is not recruited to TNFR complex I, leading to suppression of this branch of the TNF-induced signalling cascade (Heger et al., 2018).

These observations collectively suggest that LUBAC activity and linear ubiquitination of LUBAC components are tightly regulated. Yet, the posttranslational mechanisms controlling LUBAC and its inflammatory outcomes are poorly understood.

## Results

### Human HOIP is polyubiquitinated in cells

To investigate how HOIP is regulated by ubiquitination, we transiently expressed Myc-HOIP in HEK293T cells. Linear ubiquitin chains were below the detection limit in HEK293T extracts (Fig S1A), which may reflect their deubiquitination. To stabilize linear ubiquitin chains, we co-expressed the catalytically inactive mutant of OTULIN C129A, which acts as a dominant negative (Heger et al., 2018). In addition, to enrich for proteins modified with linear ubiquitin chains, we performed a pull-down using a known enrichment matrix called GST-linear-TUBE, which consists of GST fused to three tandem repeats of the linear-ubiquitin binding domain UBAN immobilized on a glutathione sepharose resin (Fig 1A) (Asaoka et al., 2016). We detected modified HOIP in the pull-down by immunoblotting (Fig S1A, lane 8, and Fig 1B, lane 1) suggesting that HOIP is ubiquitinated at least partially by linear ubiquitin chains, as reported previously with *Otulin^C129A/C129A^* knockin mouse embryonic fibroblasts (MEFs) (Heger et al., 2018).

**Figure 1.**
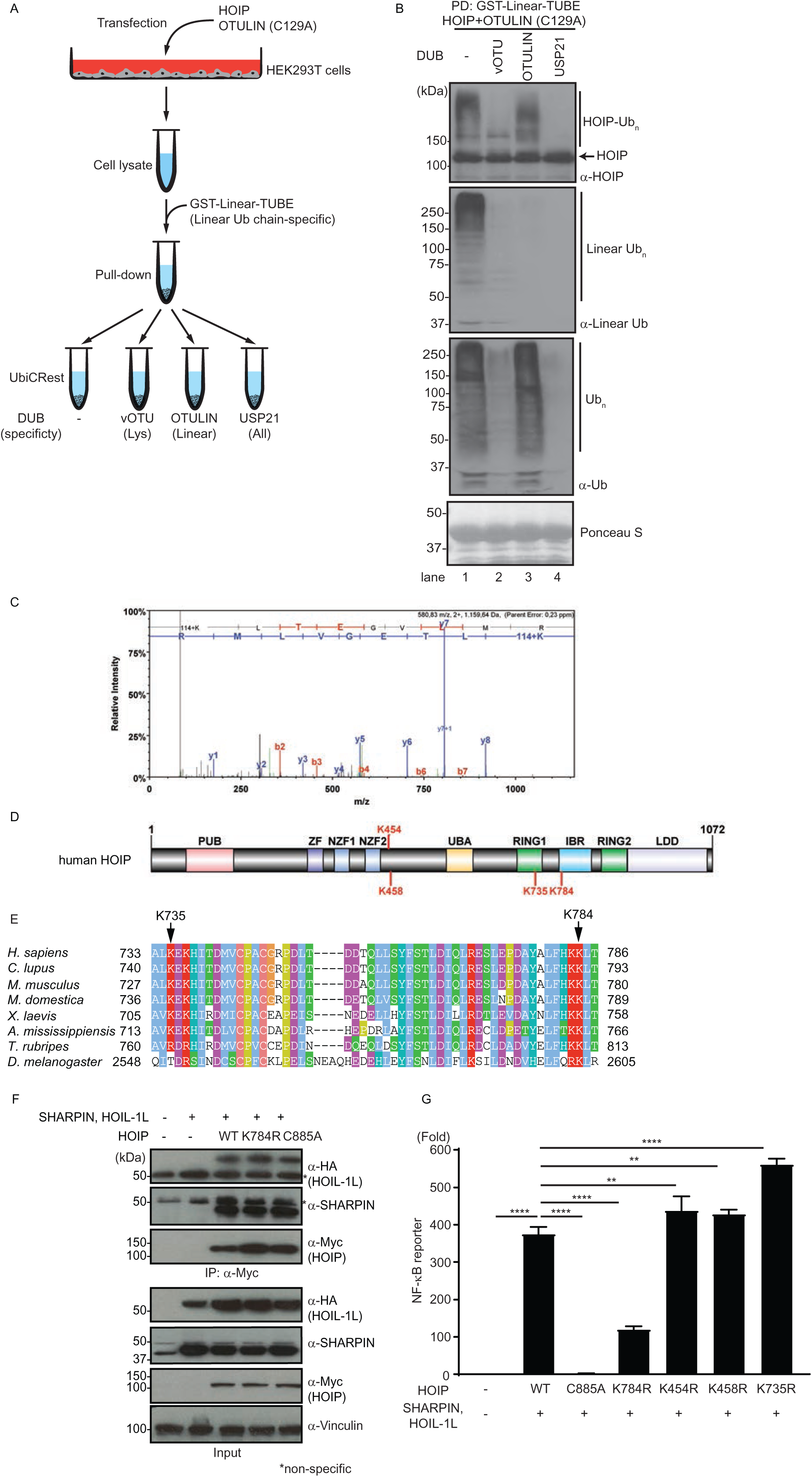
Human HOIP is ubiquitinated at K784 and regulates NF-κB. A. A scheme of procedures for UbiCRest-based assays to analyze HOIP-modification with ubiquitin chains. Total cell extracts from HEK293T cells transiently expressing Myc-HOIP and Myc-OTULIN C129A, a catalytic inactive mutant, subjected to GST-linear-TUBE pulldown followed by UbiCRest assays using recombinant deubiquitinases (vOTU, OTULIN and USP21). B. UbiCRest assays to evaluate ubiquitin chain types on HOIP examined by immunoblotting. Immunoblotting of samples using antibodies as indicated. Ponceau S staining used for monitoring GST-liner-TUBE input. Representative data shown from three independent experiments. C. Mass spectrometry spectra corresponding to a peptide containing HOIP-K784 with double Gly (114+K). D. Domains of human HOIP and identified ubiquitination sites at K454, K458, K735 and K784. E. Multiple sequence alignment of different HOIP orthologues illustrating the position K735 and K784 according to the ClustalX colour scheme. Sequences were retrieved from the NCBI protein database with the following accessions: *Homo sapiens* (NP_060469.4), *Canis lupus* (XP_005623312.1), *Mus musculus* (NP_919327.2), *Monodelphis domestica* (XP_007479924.1), *Xenopus laevis* (NP_001090429.1), *Alligator mississippiensis* (XP_006259801.1), *Takifugu rubripes* (XP_003968217.2) and *Drosophila melanogaster* (NP_723214.2). F. Co-immunoprecipitation analysis of the HOIP K784R mutant with SHARPIN and HOIL-1L using total cell extracts of HEK293T cells transiently expressing Myc-HOIP wildtype (WT), Myc-HOIP-K784R, or Myc-HOIP-C885A (a catalytic inactive mutant) with HOIL-1L-HA and Flag-SHARPIN. Anti-Vinculin antibody used to monitor the loading. Representative data shown from three independent experiments. G. Luciferase-based NF-κB gene reporter assays using Myc-HOIP wildtype (WT), a catalytic inactive mutant C885A, ubiquitination site mutants of K454R, K458R, K735R, K784R co-transfected with HOIL-1L-HA and Flag-SHARPIN. Luciferase signal was normalized to an internal Renilla control signal. Data information: In G, data are presented as mean ± SD. **p≤0.01, ****p≤0.0001 (ANOVA). n=4

To verify modification of HOIP, we used the Ubiquitin Chain Restriction (UbiCRest) method (Hospenthal et al., 2015), a DUB-based analysis (Fig 1A). The observed HOIP modification disappeared upon treatment with USP21, which hydrolyzes all linkage types of ubiquitin chains, verifying that the modification is ubiquitination (Fig 1B, lane 4). Treatment with either vOTU, which cleaves Lys-linked ubiquitin chains but not linear ubiquitin chains, or with OTULIN, which specifically cleaves linear ubiquitin chains, partially reduced the modification of HOIP (Fig 1B, lane 2 and 3). These data suggest that HOIP is polyubiquitinated with mixed linkage types of Lys and linear. OTULIN treatment diminished the levels of high-molecular weight HOIP, suggesting that linear ubiquitin chains are added on the Lys-linked ubiquitin chains.

Using mass spectrometry, we uncovered four ubiquitinated residues within human HOIP: lysine (K)454, K458, K735, K784 (Fig 1C-D, Fig S1B-D). K454 and K458 are not within any of the annotated HOIP domains and are not well-conserved (Fig 1D, Fig S1E). In contrast, K784 is within the “In Between Ring fingers” (IBR) domain, and K735 is located within the “Really Interesting New Gene” (RING1) domain (Fig 1D), and both residues are conserved in a wide range of species (Fig 1E).

### HOIP K784 regulates NF-κB activation without affecting LUBAC complex formation in cells

To evaluate the functional role of these ubiquitination sites in HOIP, we generated HOIP K454R, K458R, K735R and K784R mutants. Given that LUBAC is a regulator of NF-κB signaling, we examined these HOIP mutants using standard NF-κB reporter assays in which luciferase expression is under the control of NF-κB-response elements. Transfected cells expressed similar levels of wild type (WT) HOIP, the ubiquitination-site mutants (K454R, K458R, K735R, K784R), or a negative control (catalytically inactive mutant, C885A) (Fig 1F, S1F). As expected, we observed an increased luciferase signal in cells that co-express SHARPIN and HOIL-1L with HOIP WT, but not with HOIP C885A (Fig 1G). The luciferase signal was also significantly reduced in cells co-expressing HOIP K784R, whereas it was significantly increased, albeit mildly, in cells co-expressing HOIP K454R, K458R or K735R (Fig 1G). We observed similar results in assays without HOIL-1L or SHARPIN (Fig S1G and H).

We chose to pursue the HOIP K784 site, given that it is conserved and promotes NF-κB signaling. To investigate how linear ubiquitination at K784 affects LUBAC formation, we transiently co-expressed HOIP WT, HOIP C885A or HOIP K784R with HOIL-1L and SHARPIN in HEK293T cells, and analyzed interactions by co-immunoprecipitation. HOIP WT, HOIP K784R and HOIP C885A each interacted with HOIL-1L and SHARPIN (Fig 1F), suggesting that HOIP K784R supports LUBAC complex formation in cells. These data suggest that HOIP K784R reduces NF-κB reporter activity without compromising LUBAC complex formation.

### HOIP K784R generates unanchored linear ubiquitin chains and ubiquitinates NEMO

According to crystal structure analysis, HOIP K784 is on the surface of an alpha helix in the IBR domain, not in contact with E2 (UbcH5) or ubiquitin loaded on E2, and distant from the active site C885 (Lechtenberg et al., 2016). To determine how mutations in HOIP affect its activity, we purified recombinant HOIP proteins and performed *in vitro* ubiquitination assays. As expected, the HOIP C885A catalytic mutant did not generate unanchored linear ubiquitin chains nor did it ubiquitinate NEMO (Fig 2A, Fig S2A). Further, HOIP C885A was not polyubiquitinated (Fig 2A, Fig S2A), suggesting that HOIP is modified dependently on its own catalytic activity *in vitro*. Both HOIP WT and HOIP K784R generated unanchored linear ubiquitin chains and ubiquitinated the LUBAC substrate NEMO, when co-incubated with SHARPIN and HOIL-1L (Fig 2A). In the absence of SHARPIN, the ubiquitination signal was substantially reduced in reactions with HOIP K784R compared to HOIP WT (Fig S2A). These data suggest that the HOIP K784R mutant, in a complex with both HOIL-1L and SHARPIN, can ubiquitinate substrates *in vitro*, though with altered kinetics compared to HOIP WT.

**Figure 2.**
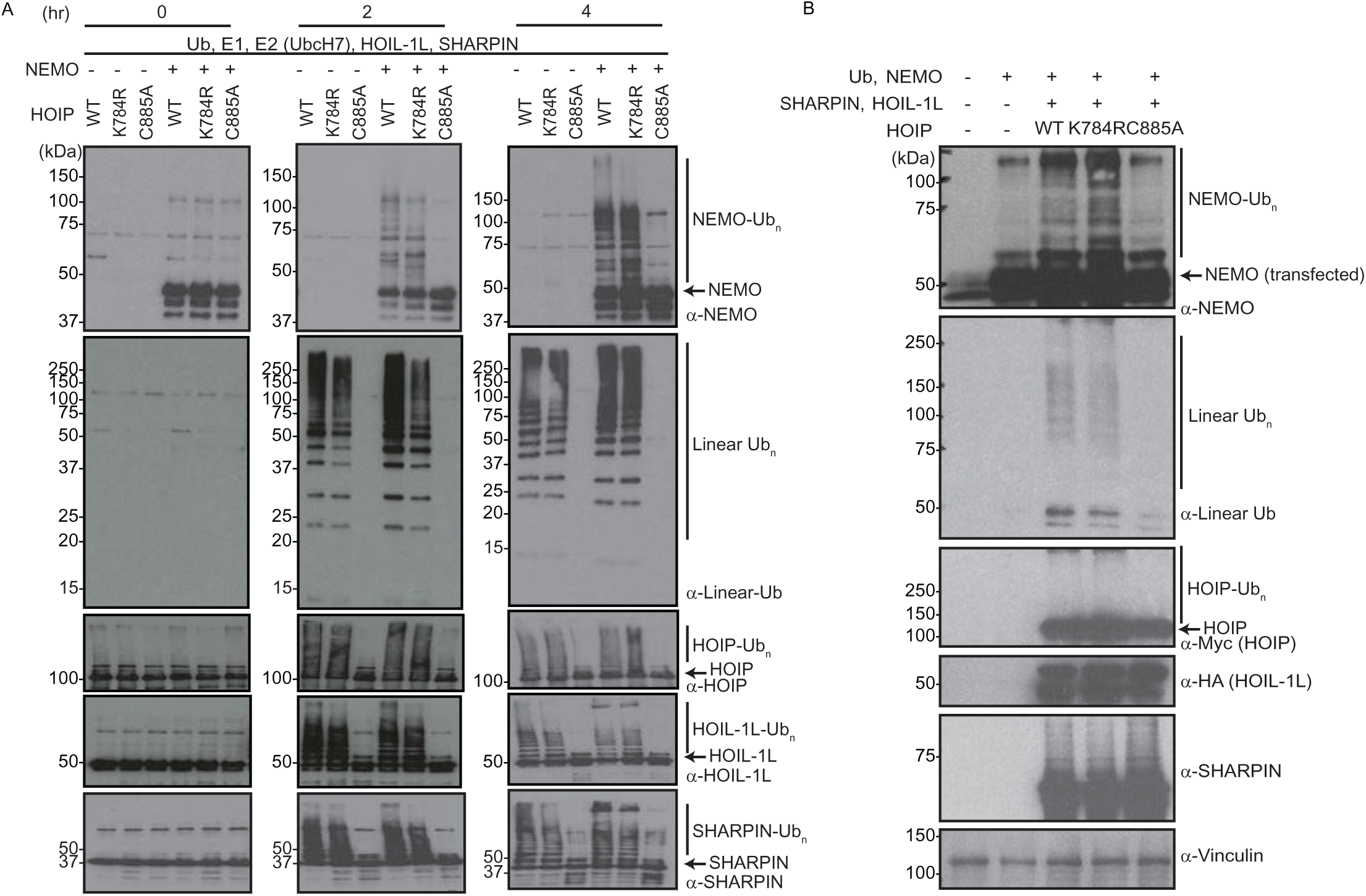
The human HOIP K784R mutant as a part of the LUBAC generates linear ubiquitin chains and ubiquitinates its substrate NEMO *in vitro* and in cells. A. *In vitro* ubiquitination assays for the indicated times using the recombinant proteins of ubiquitin (Ub), E1, E2 (UbcH7), HOIP (WT, K784R or C885A mutant), HOIL-1L, SHARPIN and NEMO. Immunoblotting of Linear ubiquitin chains, NEMO, HOIP, HOIL-1L and SHARPIN detected by using antibodies as indicated. Representative data from three independent experiments. B. Immunoblotting to detect ubiquitination of NEMO in HEK293T cells transiently expressing Flag-NEMO, GFP-SHARPIN, and HOIL-1L-HA with Myc-HOIP wildtype (WT), Myc-HOIP K784R or Myc-HOIP C885A. Total cell lysates in denaturing conditions subjected to SDS-PAGE followed up by immunoblotting using antibodies as indicated. Anti-Vinculin antibody to monitor loading. Representative data from three independent experiments.

To test HOIP activity in cells, we transiently expressed HOIP WT, HOIP K784R or HOIP C885A with HOIL-1L and SHARPIN in HEK293T cells. Cells expressing HOIP WT and HOIP K784R displayed similar levels of linear ubiquitin chains and polyubiquitinated NEMO (Fig 2B), whereas cells expressing HOIP C885A lacked both ubiquitination events. These results collectively indicate that HOIP K784R, as a part of LUBAC, can ubiquitinate NEMO *in vitro* and in cells. The observations of HOIP ubiquitination dependent on its catalytic site C885 (Fig 2A and B) and HOIP ubiquitination by mixed linkage types of chains in cells (Fig 1B) suggest that HOIP ubiquitination is at least partially self-ubiquitination in cells.

### *Hoip^K778R/K778R^* mice do not have overt developmental defects

Given that human HOIP K784R disrupted NF-κB signaling in the cellular reporter cell assay without substantially affecting HOIP catalytic activity or LUBAC formation in HEK293T cells, we investigated the endogenous function of HOIP K784 *in vivo*. We used CRISPR-Cas9 to generate homozygous knockin mice with a substitution at HOIP K778, the equivalent residue to K784 in mice (*Hoip^K778R/K778R^*, Fig S3A and S3B). *Hoip^K778R/K778R^* mice were born at the nearly expected ratio from crosses of *Hoip^+/K778R^* mice and displayed no obvious developmental phenotypes (Fig 3A and B). These observations are in contrast to HOIP loss-of-function mice, which are embryonic lethal (Peltzer et al., 2014) (Emmerich et al., 2013) (Hrdinka & Gyrd-Hansen, 2017).

**Figure 3.**
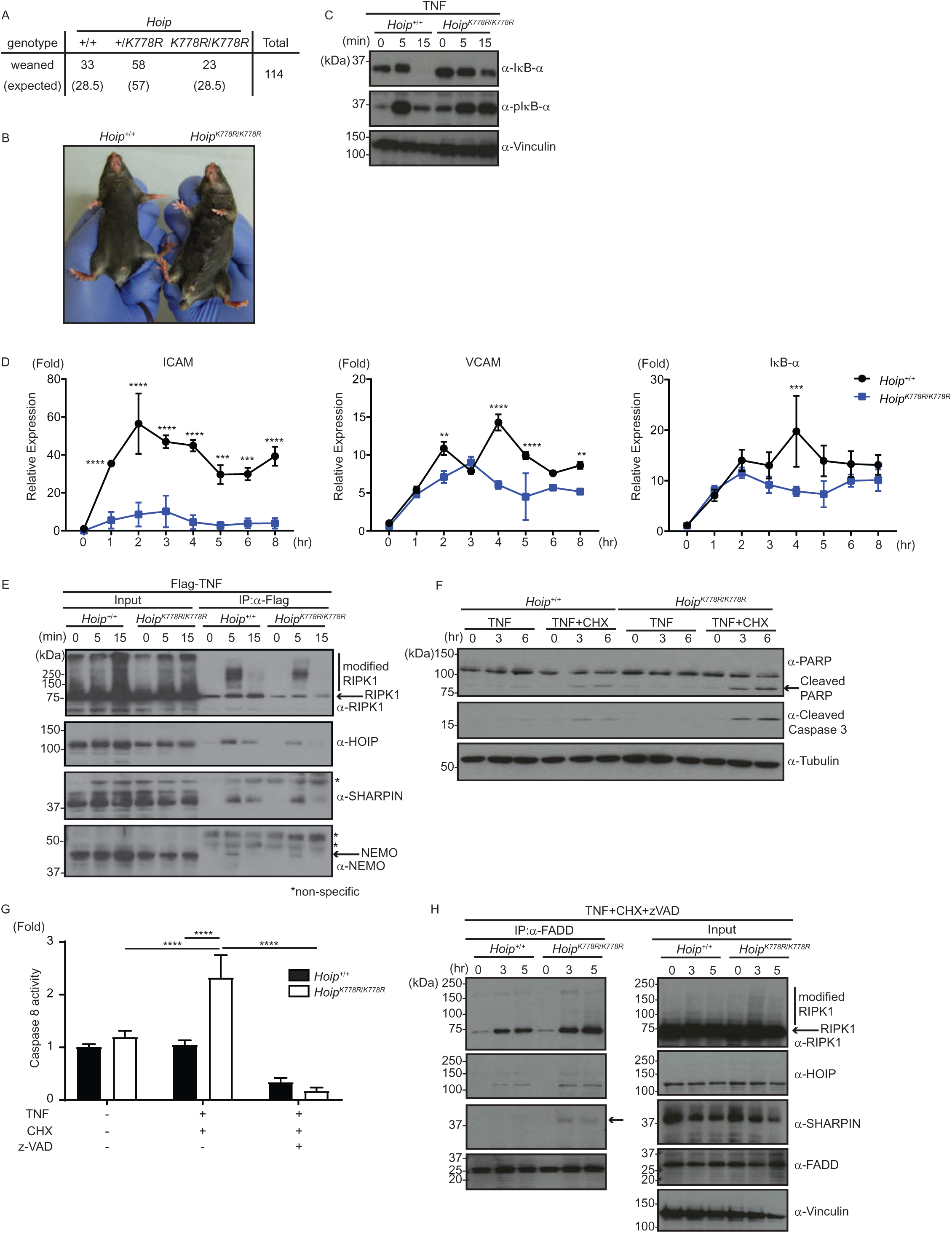
No obvious developmental defect in *Hoip^K778R/K778R^* knockin mice is observed while TNF-responses are suppressed in *Hoip^K778R/K778R^* cells. A. Numbers of weaned mice of the indicated genotypes from *Hoip^+/K778R^*; *Sharpin^+/cpdm^* crosses. B. A gross appearance image of *Hoip^+/+^* and *Hoip^K778R/K778R^* male mice at 6-week old. C. Immunoblotting to detect TNF-induced degradation and phosphorylation of IκB-α in immortalized *Hoip^+/+^* and *Hoip^K778R/K778R^* MEFs treated with human TNF (20ng/ml) for the indicated times. Anti-Vinculin antibody for monitoring loading amount. Representative data from three independent experiments. D. Induction of TNF-dependent NF-κB target genes, ICAM, VCAM and IκB-α in *Hoip^+/+^* or *Hoip^K778R/K778R^* immortalized MEFs determined by qRT-PCR. RNA extraction and cDNA synthesis from MEFs treated with hTNF (20ng/ml) for the indicated time subjected to examine transcripts of ICAM, VCAM and IκB-α. Normalization to β-actin. E. Immunoblotting of TNF-immunoprecipitation to examine TNFR complex I formation in immortalized *Hoip^+/+^* or *Hoip^K778R/K778R^* MEFs treated with Flag-hTNF (1µg/ml) for the indicated time using indicated antibodies. Representative data from three independent experiments. F. TNF-dependent induction of cleavage of PARP and Caspase 3 in primary *Hoip^+/+^* or *Hoip^K778R/K778R^* MEFs determined by immunoblotting. Total cell extracts of MEFs treated with hTNF (100ng/ml) and CHX (1µg/ml) for the indicated times subjected to SDS-PAGE followed by immunoblotting using antibodies as indicated. Anti-α-Tubulin antibody used to monitor loading amount. Representative data of two independent experiments. G. TNF-induced Caspase 8 activity in *Hoip^+/+^* or *Hoip^K778R/K778R^* immortalized MEFs treated with hTNF (100ng/ml) with or without Cycloheximide (CHX) (1µg/ml) or z-VAD (20µM). H TNF-receptor complex II formation in *Hoip^+/+^* or *Hoip^K778R/K778R^* immortalized MEFs. Total cell extracts of MEFs treated with hTNF (100ng/ml), CHX (1µg/ml) and zVAD (25µM) for the indicated timeimmunoprecipitated using an anti-FADD antibody. Recruitment of RIPK1, HOIP and SHARPIN monitored by immunoblotting. Representative of two independent experiments. Data information: In (D and G), data are presented as mean ± SD. **p≤0.01, ***p≤0.001, ****p≤0.0001 (ANOVA, n=3 in D and n=4 in G).

### TNF-induced NF-κB activation is suppressed in *Hoip^K778R/K778R^* MEFs

To analyze if HOIP K778 is involved in the regulation of TNF-dependent NF-κB signaling, we derived MEF lines from *Hoip^+/+^* and *Hoip^K778R/K778R^* mice and stimulated them with TNF (Fig 3C and D, Fig S3C). We found that TNF-induced phosphorylation of IκB-α is prolonged and degradation of IκB-α was reduced in *Hoip^K778R/K778R^* MEFs compared with *Hoip^+/+^* MEFs (Fig 3C). Furthermore, the TNF-induced transcription of some NF-κB target genes, such as ICAM, VCAM and IκB-α, was significantly reduced in *Hoip^K778R/K778R^* MEFs compared to *Hoip^+/+^* MEFs (Fig 3D), whereas TNF-induced gene induction of A20 was unaffected (Fig S3C).

To elucidate the step of TNF-dependent signaling that is affected in *Hoip^K778R/K778R^* MEFs, we examined the formation of TNFR complex I. Upon TNF-treatment, RIPK1, HOIP, SHARPIN, and NEMO co-immunoprecipitated with TNF in both *Hoip^+/+^* and *Hoip^K778R/K778R^* MEFs (Fig 3E), indicating recruitment to TNFR complex I. However, recruitment of RIPK1, HOIP and SHARPIN were mildly decreased in *Hoip^K778R/K778R^* MEFs compared to WT MEFs (Fig 3E).

Collectively, these data indicate that HOIP K778R significantly suppresses the TNF-induced NF-κB signaling cascade in MEFs, coincident with slightly diminished formation of TNFR complex 1.

### *Hoip^K778R/778KR^* MEFs are sensitized to apoptosis

LUBAC plays a role in the anti-apoptotic branch of the TNF pathway (Asaoka & Ikeda, 2015, Sasaki & Iwai, 2015, Walczak, 2011). Therefore, we assessed the ability of *Hoip^K778R/K778R^* MEFs to resist TNF-dependent cell death. To this end, we examined TNF-mediated induction of the active form of Caspase 3 (cleaved-Caspase 3), which is a so-called apoptosis executioner caspase, and cleavage of its substrate, PARP (Fig 3F). We also treated cells with cycloheximide (CHX), an inhibitor of translation which sensitizes cells to TNF-induced apoptosis (Kumari et al., 2014, Rahighi et al., 2009). Compared to *Hoip^+/+^* MEFs, *Hoip^K778R/K778R^* MEFs displayed elevated levels of both cleaved-Caspase 3 and cleaved PARP after treatment with TNF and CHX (Fig 3F).

We also measured the activity of an apoptosis initiator caspase, Caspase 8, using luminescent assays in *Hoip^+/+^* and *Hoip^K778R/K778R^* primary MEFs. We observed significantly higher levels of Caspase 8 activity in *Hoip^K778R/K778R^* MEFs than in *Hoip^+/+^* MEFs after treatment with TNF and CHX (Fig 3G). Similar responses were observed in MEFs treated with a different cell death ligand, Fas-ligand (FasL) (Fig S3D). As expected, treatment with the pan-caspase inhibitor z-VAD eliminated Caspase 8 activity (Fig 3G, Fig S3D).

TNF- and FasL-induced apoptosis pathways are mediated through the TNFR complex II and the death-inducing signaling complex (DISC), respectively. These signaling pathways have overlapping components including FADD and Caspase 8. Thus, we hypothesized that HOIP K784 ubiquitination regulates those cell death-inducing complexes. To address this point, we examined TNFR complex II formation in *Hoip^+/+^* and *Hoip^K778R/K778R^* MEFs. We treated MEFs with TNF, CHX and z-VAD and immunoprecipitated the TNFR complex II component FADD. We observed an enhanced complex formation between FADD and RIPK1, HOIP and SHARPIN in *Hoip^K778R/K778R^* MEFs compared to *Hoip^+/+^* MEFs. Thus, formation of the TNFR complex II in response to TNF and CHX is elevated in *Hoip^K778R/K778R^* MEFs compared to *Hoip^+/+^* MEFs.

### *Hoip^K778R/K778R^; Sharpin ^cpdm/cpdm^* mice display TNFR1-dependent embryonic lethality

Given that *Hoip^K778R/K778R^* MEFs are sensitized to TNF-induced apoptosis, and that the LUBAC component SHARPIN plays a role in the same apoptosis pathway (Ikeda et al., 2011), we investigated a potential cooperation between HOIP and SHARPIN *in vivo*. To this end, we attempted to generate *Hoip^K778R/K778R^* mice in the *Sharpin^cpdm/cpdm^* background, which do not express functional SHARPIN protein. We did not observe any *Hoip^K778R/K778R^*;*Sharpin^cpdm/cpdm^* newborn mice from a cross of *Hoip^+/K778R^*;*Sharpin^+/cpdm^* mice (Fig 4A), suggesting embryonic lethality. To examine the embryonic development of *Hoip^K778R/K778R^*;*Sharpin^cpdm/cpdm^* mice, we crossed *Hoip^K778R/K778R^*;*Sharpin^+/cpdm^* mice and analyzed embryos at E12.5 and E13.5 (Fig 4B and C, Fig S4A). We observed embryos of all genotypes at both these stages (Fig 4B). However, *Hoip^K778R/K778R^*;*Sharpin^cpdm/cpdm^* embryos were found to be unhealthy with grossly evident regions of hemorrhage, suggesting the possibility of lethality at, or immediately subsequent to, these stages (Fig S4A, Fig 4C).

**Figure 4.**
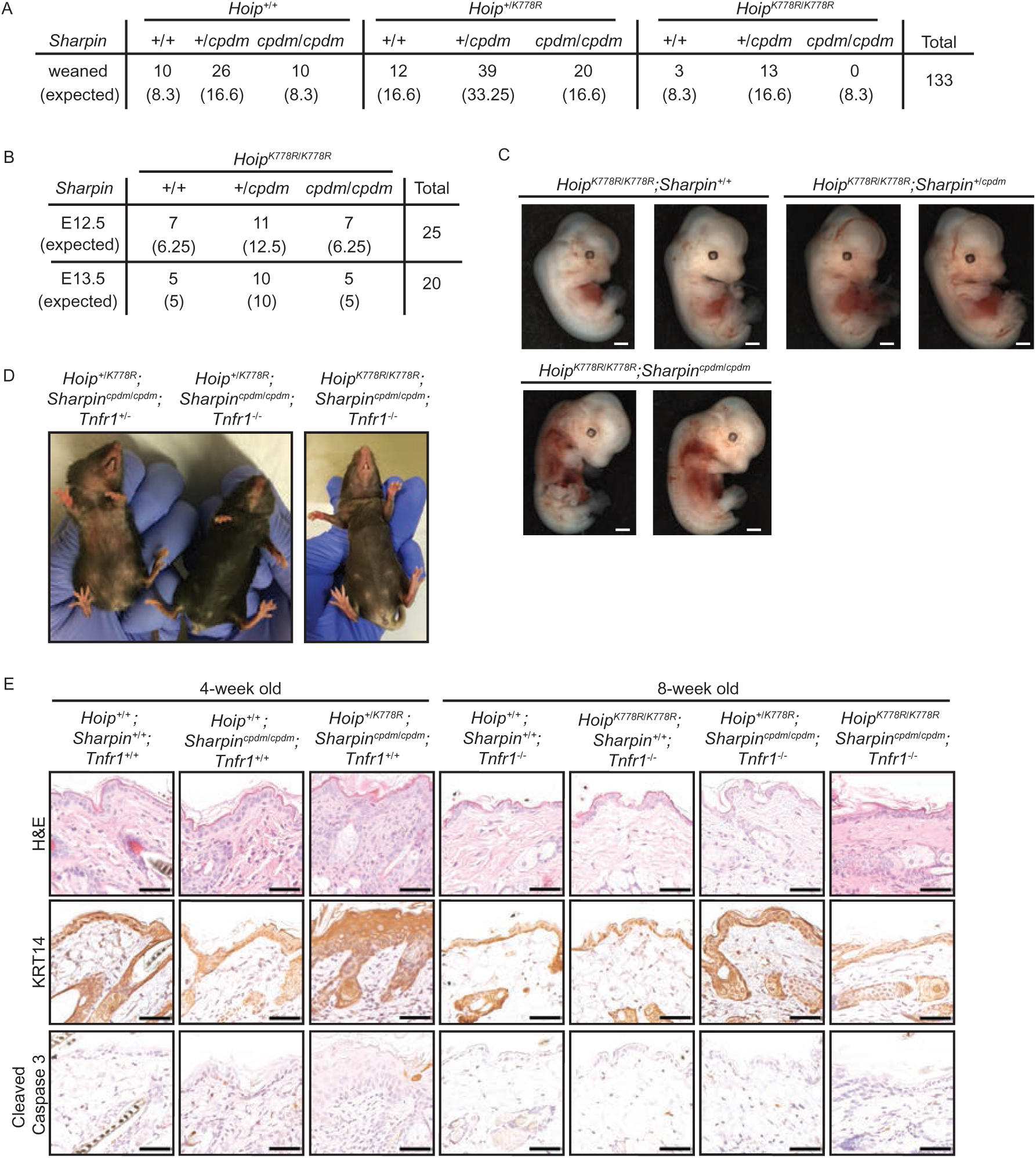
SHARPIN-deficiency leads *Hoip^K778R/K778R^* mice to embryonic lethality, which is rescued by TNFR1 knockout. A. Numbers of weaned mice of the indicated genotypes from crosses of *Hoip^+/K778R^; Sharpin^+/cpdm^* mice. B. Number of the embryos of the indicated genotype at E12.5 and E13.5 from crosses of *_Hoip_K778R/K778R_; Sharpin_+/cpdm* _mice._ C. Gross appearance images of *Hoip^K778R/K778R^; Sharpin^+/+^*, *Hoip^K778R/K778R^; Sharpin^+/cpdm^* and *Hoip^K778R/K778R^; Sharpin^cpdm/cpdm^* embryos at E13.5. Representative pictures from 7 embryos each. Scale bars:1mm D. Gross appearance images of *Hoip^+/K778R^;Sharpin^cpdm/cpdm^;Tnfr1^+/-^* and *Hoip^K778R/+^; Sharpin^cpdm/cpdm^; Tnfr1^−/−^* female mice at 6 weeks of age, and a female *Hoip^K778R/K778R^; Sharpin^cpdm/cpdm^; Tnfr1^−/−^* mouse at 8 weeks of age. E. H&E staining and Keratin 14 (KRT14) immunostaining of dorsal skin sections of the indicated genotypes at 4-week or 8-week old of age. Scale bars: 50um.

*Sharpin^cpdm/cpdm^* mice display a systemic inflammatory phenotype that requires TNFR1 (Kumari et al., 2014, Rickard et al., 2014). To elucidate if the embryonic lethality of *Hoip^K778R/K778R^*;*Sharpin^cpdm/cpdm^* mice also depends on TNFR1, we tested whether it is rescued by TNFR1 knockout. Strikingly, the embryonic lethality is rescued in *Hoip^K778R/K778R^*;*Sharpin^cpdm/cpdm^*;*Tnfr1^−/−^* mice (Fig 4D), and skin inflammation was nearly absent until at least 8 weeks after birth (Fig 4E). Furthermore, qualitative reduction of inflammation in tissues such as the lung and liver was also observed in 8-week old *Hoip^K778R/K778R^*;*Sharpin^cpdm/cpdm^*;*Tnfr1^−/−^* mice (Fig S4B).

### *Hoip^+/K778R^; Sharpin^cpdm/cpdm^* mice show early onset of severe proliferative dermatitis with keratinocyte apoptosis

Remarkably, heterozygosity of *Hoip^+/K778R^* was sufficient to bring about an early onset of dermatitis in *Sharpin^cpdm/cpdm^* mice (Fig 4D and E, Fig 5A and B). At 4 weeks of age, these mice developed chronic proliferative dermatitis characterized by acanthosis, hyperkeratosis, dermal inflammatory cell infiltrates and keratinocyte apoptosis (detected by cleaved Caspase 3). The acanthosis was further confirmed by the thickened Keratin 14 (KRT14) positive zone in the epidermis (Fig 4E). In contrast, *Hoip^+/+^;Sharpin^cpdm/cpdm^* mice showed no clear sign of skin lesions at this age (Fig 5A and B). The extent of the chronic proliferative dermatitis in *Hoip^+/K778R^*;*Sharpin^cpdm/cpdm^* mice was further quantified by measurements of total epidermal thickness, keratin layer thickness and squamous epithelial layer thickness (Fig5 C-E). Increased dermal inflammatory cell infiltration in *Hoip^+/K778R^*;*Sharpin^cpdm/cpdm^* mice was further confirmed by the macrophage marker, F4/80, and the monocyte/granulocyte/neutrophil maker Ly6G (Fig 5B). Importantly, apoptotic epidermal keratinocytes with activated Caspase 3 positivity were also prominent in these skin sections (Fig 5B, Cleaved Caspase 3 panels). These features resemble those of *Sharpin^cpdm/cpdm^* mice at an older age (8-week old). In addition to the skin, *Hoip^+/K778R^*;*Sharpin^cpdm/cpdm^* mice showed a multi-systemic inflammatory phenotype with immune cell infiltrates in visceral organs such as the lung and liver (Fig S5A). As in *Sharpin^cpdm/cpdm^* mice, small intestinal Peyer’s patches, secondary lymphoid structures, were notably absent in *Hoip^+/K778R^*;*Sharpin^cpdm/cpdm^* mice (Fig S5A). In contrast to the enlarged spleens in *Sharpin^cpdm/cpdm^* mice, spleens were smaller in *Hoip^+/K778R^*;*Sharpin^cpdm/cpdm^*(Fig S5B). In the spleens of *Sharpin^cpdm/cpdm^* and *Hoip^+/K778R^*;*Sharpin^cpdm/cpdm^* mice, white pulp follicular architecture was obscured in concert with enhanced myeloid hyperplasia in the red pulp (Fig S5A, S5B). The skin inflammatory phenotype of *Hoip^+/K778R^*;*Sharpin^cpdm/cpdm^* mice was mitigated by TNFR1 knockout (Fig 4D and E). These results collectively suggest that ubiquitination of HOIP at K778 in mice collaborates with SHARPIN to regulate TNF-induced inflammation and cell death.

**Figure 5.**
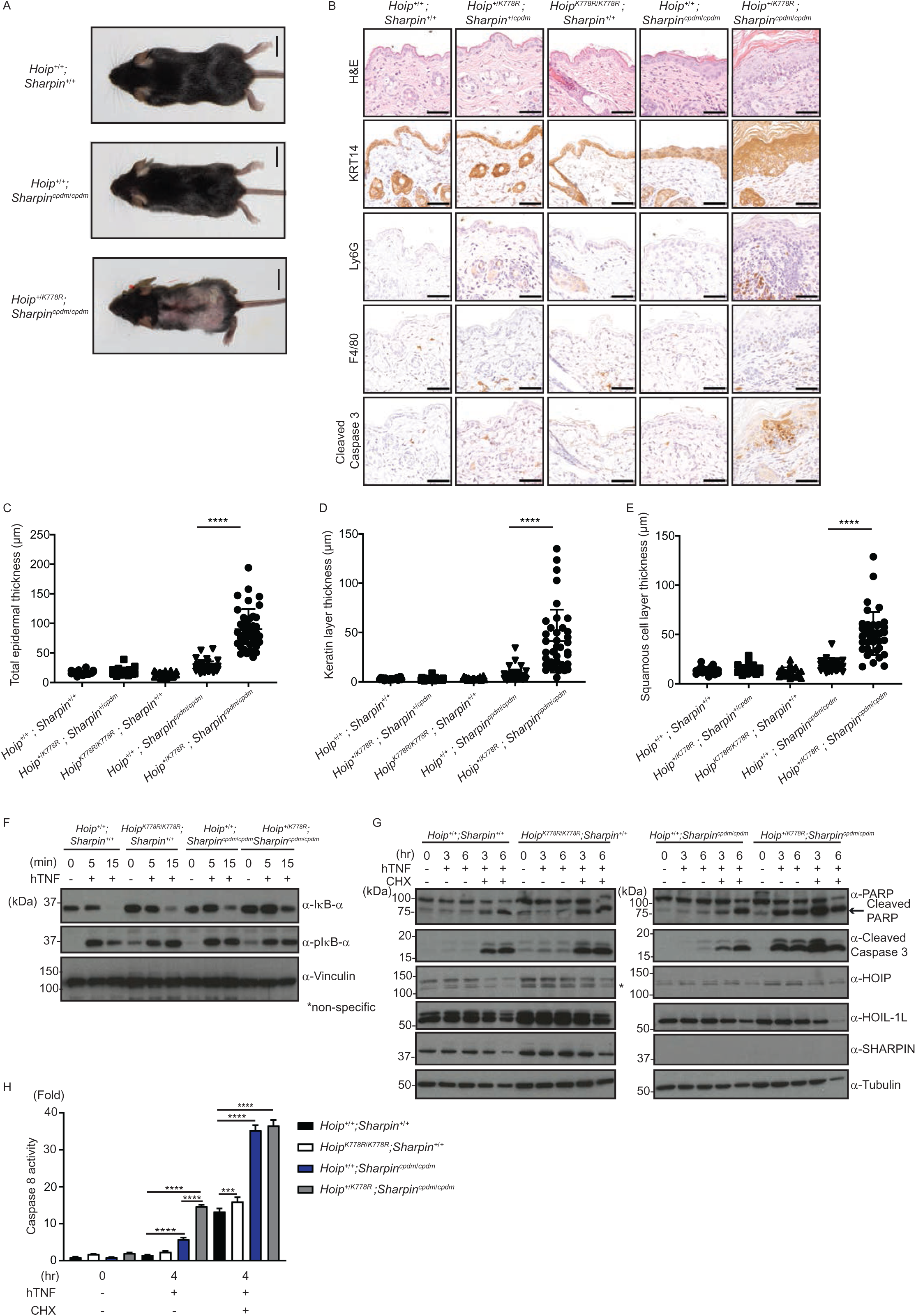
SHARPIN-deficiency in *Hoip^+/K778R^* heterozygous mice leads to early onset of skin inflammation accompanied with apoptosis induction. A. Gross appearance images of mice of the indicated genotypes (male mice at 4-week old). Scale bars: 10mm. B. Immunostaining of dorsal skin sections (H&E, Keratin 14 (KRT14), Ly6G, F4/80 and Cleaved Caspase 3) of mice with the indicated genotypes. Scale bars: 50µm. C-E. Measurements of dorsal skin sections (Total epidermis skin thickness, Keratin layer thickness and Squamous cell layer thickness) obtained from male mice of the indicated genotypes. Each dot on the scatter dot plot represents one focus point of the measurement. (N=20, 20, 30, 40, 40 for *Hoip^+/+^; Sharpin^+/+^*, *Hoip^K778R/K778R^; Sharpin^+/+^, Hoip^+/K778R^; Sharpin^+/cpdm^*, *Hoip^+/+^; Sharpin^cpdm/cpdm^, Hoip^+/K778R^; Sharpin^cpdm/cpdm^*, respectively.) F. Immunoblotting to examine TNF-induced degradation and phosphorylation of IκB-α in immortalized MEFs of the indicated genotypes using total cell extracts of MEFs treated with hTNF (20ng/ml) for the indicated times. Representative of three independent experiments. G. Immunoblotting of TNF-induced cleavage of PARP and caspase 3 in immortalized MEFs of the indicated genotypes using total cell extracts of MEFs treated with hTNF (100ng/ml) with or without CHX (1µg/ml) for indicated times. Representative of three independent experiments. H. TNF-induced Caspase 8 activity in immortalized MEFs of indicated genotype. Luminol-dependent activity of Caspase 8 in immortalized MEFs treated with hTNF (100ng/ml) with or without CHX (1µg/ml) or z-VAD (20µM). Data information: In (C-E and H), data are presented as mean ± SD. (n=4 for H, ***p≤0.001, ****p≤0.0001, ANOVA).

To test whether the TNFR1-dependent inflammation and apoptosis phenotypes in *Hoip^+/K778R^*;*Sharpin^cpdm/cpdm^* mice are cell autonomous, we established immortalized MEFs and examined TNF-induced signaling cascades, including NF-κB activation and apoptosis (Fig 5F-H). TNF-induced degradation of IκB-α was further delayed in *Hoip^+/K778R^*;*Sharpin^cpdm/cpdm^* MEFs, similar to *Hoip^K778R/K778R^* MEFs and *Sharpin^cpdm/cpdm^* MEFs (Fig 5F). Cleaved PARP, cleaved Caspase 3 and activated Caspase 8 were elevated in *Hoip^+/K778R^*;*Sharpin^cpdm/cpdm^* MEFs compared to *Hoip^+/+^* MEFs, *Hoip^K778R/K778R^* MEFs and *Sharpin^cpdm/cpdm^* MEFs treated with TNF, particularly without CHX (Fig 5G and H). We observed similar results with primary ear-derived fibroblasts (Fig S5C). These results indicate that HOIP ubiquitination at K778 cooperates with SHARPIN to promote TNF-induced cell survival in a cell autonomous manner.

## Discussion

Our findings demonstrate that a site-specific ubiquitin modification of HOIP (K784 in human, K778 in mouse) cooperates with SHARPIN to impact TNF-dependent signaling cascades and immune responses in mice (Fig 6). Importantly, HOIP K784R is still ubiquitinated at other sites, both in cells and *in vitro*, emphasizing a specific requirement for ubiquitination at K784. TNFR1 knockout rescues the mouse phenotypes arising from HOIP K778R and SHARPIN-deficiency, indicating the dependency on TNFR1 signalling. The K-to-R substitution in HOIP (K784R in human or K778R in mouse) did not alter HOIP levels, indicating that ubiquitination at this site does not regulate proteasomal degradation.

**Figure 6.**
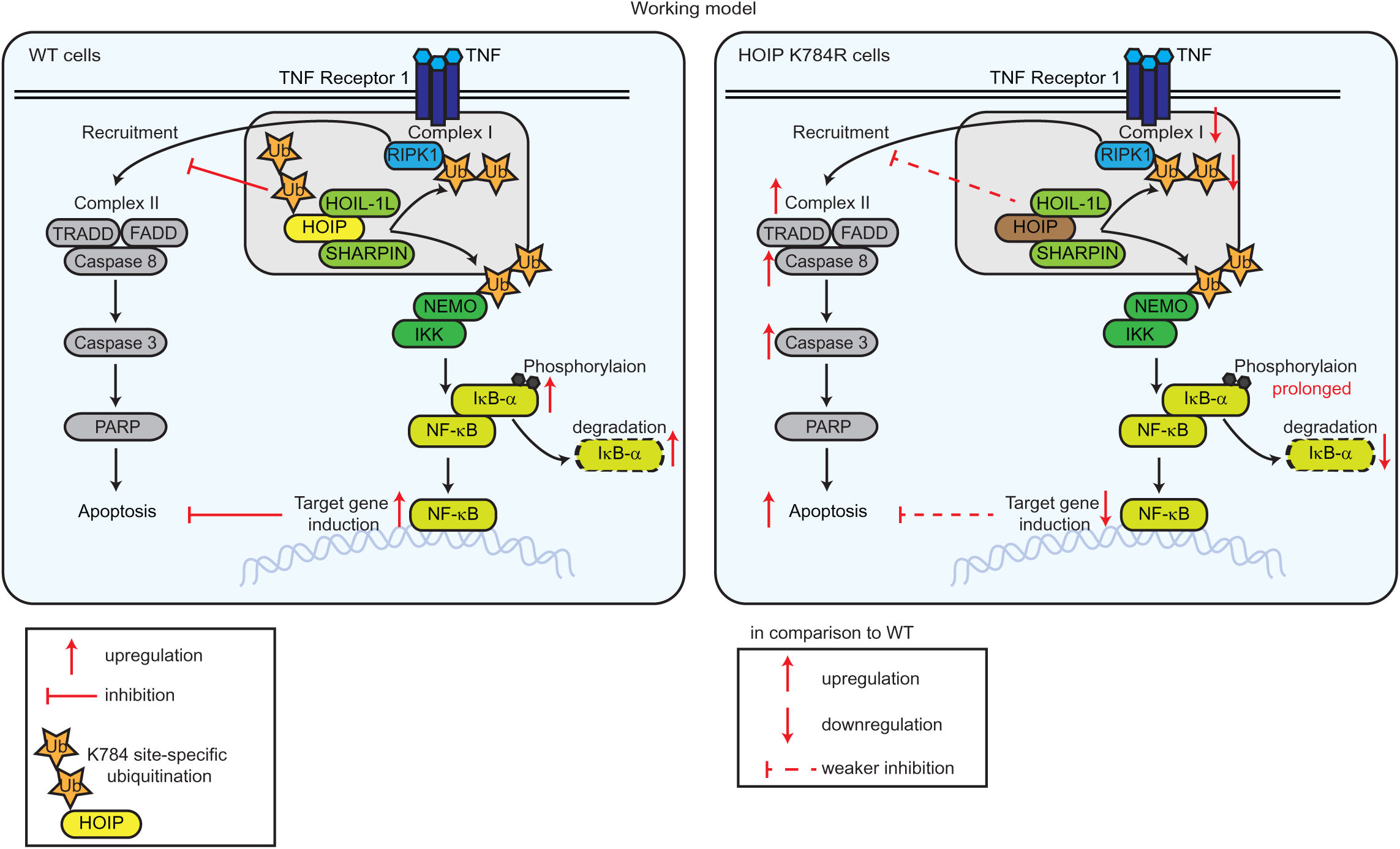
A proposed model of HOIP-site specific ubiquitination in the regulation of the TNF signaling cascades. A working model describing a role of site-specific ubiquitination of HOIP in the regulation of the TNF-induced NF-κB and apoptosis pathways based on this study. Schematics of simplified TNF signaling in wild type cells (WT, left panel) and HOIP-K784R cells (right panel) are shown. Upon TNF-binding to TNF receptor 1, complex I formation, which includes RIPK1 and LUBAC components (HOIP, HOIL-1L and SHARPIN) is formed. In this signaling cascade, LUBAC ubiquitinates NEMO and RIPK1. Site specific ubiquitination of HOIP at K784 plays a role in inducing TNF-dependent NF-κB target genes and in inhibition of apoptosis by inhibiting RIPK1 recruitment into TNF receptor complex II.

*Hoip^K778R/K778R^* knockin mice, generated in this study, manifest no overt phenotype for at least 96 weeks after birth. Consistent with our NF-κB gene reporter assays using HOIP K784R, we found that TNF-induced NF-κB activation was suppressed in *Hoip^K778R/K778R^* cells compared to *Hoip^+/+^* cells. Since the recruitments of HOIP, SHARPIN and modified and unmodified-RIPK1 to TNFR complex I were mildly reduced, we speculate that this site-specific ubiquitination of HOIP might regulate TNFR complex I formation in cells. Apoptosis induced by TNF and CHX was clearly enhanced in *Hoip^K778R/K778R^* cells. FADD showed enhanced interactions with RIPK1, HOIP and SHARPIN in the TNFR complex II in *Hoip^K778R/K778R^* cells relative to wild type cells, after treatment with TNF and CHX. These data suggest that ubiquitination of HOIP at K778 (in mice) plays a critical role in the apoptosis pathway by regulating the formation of downstream signalling complexes.

Embryonic lethality was observed in *Hoip^K778R/K778R^* knockin mice with concurrent loss of SHARPIN. Additionally, regions of hemorrhage were grossly evident in *Hoip^K778R/K778R^*;*Sharpin^cpdm/cpdm^* embryos at the E12.5-E13.5 stages. HOIP and HOIL-1L are critical for embryonic development in mice, and mouse knockouts of these LUBAC components result in hyper-induction of apoptosis in embryos (Peltzer et al., 2018, Peltzer et al., 2014). Strikingly, a heterozygous allele of *Hoip^+/K778R^* is sufficient to accelerate skin inflammation and apoptosis in SHARPIN-deficient mice. These data indicate collaborative roles of SHARPIN and HOIP site-specific ubiquitination at K784.

Based on the known co-crystal structure of the HOIP-C-terminal fragment with ubiquitin-loaded E2 (UbcH5), HOIP K784 within the IBR domain does not directly contact the ubiquitin or E2 (Lechtenberg et al., 2016). Furthermore, our *in vitro* ubiquitination assays using recombinant LUBAC components suggest that the suppressed activity of the NF-κB gene reporter is not due to substantial loss of LUBAC activity. However, *in vitro* reactions lacking SHARPIN show that linear ubiquitination is reduced with HOIP K784R relative to HOIP WT, which could in part explain why HOIP K784R knockin mice with no apparent phenotype become embryonic lethal with SHARPIN-deficiency. We observed complete loss of ubiquitination of HOIP C885A relative to HOIP WT, indicating that HOIP ubiquitination relies on its catalytic activity. However, the E3 ligases that directly ubiquitinate HOIP in the TNF signalling cascade in cells is not known. HOIP is modified by mixed-linkage types of linear and Lys-linked ubiquitin chains in cells, suggesting that additional E3 ligases are involved.

A method to generate a ubiquitination mimic of the substrate to directly address the impact of site-specific ubiquitination, especially by ‘poly’ubiquitination is not yet established. Thus, we mutated HOIP K784 to abolish ubiquitination at this site. Although any mutation can yield non-specific negative effects, HOIP K784R retained catalytic activity but altered the TNF-induced inflammatory by diminishing TNFR complex I but enhancing TNFR complex II formation and function. Thus, the effects we observed with HOIP K784R mutant in cell signalling are not due to non-specifics on protein folding.

Deubiquitinases are known to regulate HOIP-ubiquitination. In a previous study, expression of inactive OTULIN C129A mutant in cells leads to hyper-ubiquitination of LUBAC components, including HOIP, and prevents proper activation of TNF-induced signalling (Heger et al., 2018). We now show that site-specific ubiquitination of HOIP influenced TNFR complex formation. As a next step, it would be important to understand how the balance of OTULIN and LUBAC is controlled in different cell types to regulate immune responses.

In conclusion, our study has uncovered a new type of regulation of the ubiquitin ligase HOIP by site-specific ubiquitination, which is balances inflammatory responses by promoting the formation of TNFR complex I and inhibiting the formation of TNFR complex II. Similar to the regulation of kinases by phosphorylation, site-specific ubiquitin modification of ubiquitin ligases might regulate their activity and function in biology.

## Materials and methods

### Plasmids

pBABE-puro-Flag-human SHARPIN, pEGFP-C1-human SHARPIN, pGEX-6P-1-human HOIP, pGEX-4T-1-Linear TUBE, pGEX-6P-1-human OTULIN (WT and C129A), and pcDNA3-human-Ubiquitin were previously described (Asaoka et al., 2016, Ikeda et al., 2011). pGEX-6P-1-human NEMO, pGEX-6P-1-human HOIL-1L, pGEX-6P-1-human SHARPIN and pRK5-Myc-human OTULIN (WT and C129A) were cloned using a standard subcloning method. All the point mutants in pcDNA3-Myc-human HOIP (K454R, K458R, K735R, K784R and C885A) and pGEX-6P-1-human HOIP (K784R and C885A) were generated by site directed mutagenesis. Sequences of all the plasmids generated for this study were confirmed by Sanger sequencing. pcDNA3-Myc-human HOIP and pcDNA3-human-HOIL-1L-HA were from Kazuhiro Iwai (Tokunaga et al., 2009), pGex6P-1-human UbcH7, pET49b-human HOIP UBA-RBR-C (aa 476-1072) and pET49b-human HOIL-1L (C460A) were from Katrin Rittinger (Stieglitz et al., 2012). pOPINK-vOTU (CCHFV OTU, aa1-183) (Addgene plasmid #61589) (Akutsu et al., 2011) and pOPINS-USP21 (USP, aa 196-565) (Addgene plasmid #61585)(Ye et al., 2011) were gifts from David Komander.

Primer sequences used for the site-directed mutagenesis were the following.

HOIP K454R: forward primer 5’-GCCAGCTCTTTGGAAAGGGGACCCCCCAAG-3’, reverse primer 5’-CTTGGGGGGTCCCCTTTCCAAAGAGCTGGC-3’, HOIP K458R: forward primer 5’-GAAAAGGGACCCCCCAGGCCTGGGCCCCCA-3’, reverse primer 5’-TGGGGGCCCAGGCCTGGGGGGTCCCTTTTC-3’, HOIP K735R: forward primer: CACTTCACCATCGCCTTGAGGGAGAAGCACATC-3’, reverse primer 5’-GATGTGCTTCTCCCTCAAGGCGATGGTGAAGTG-3’, HOIP K784R forward primer 5’-GCGTTGTTCCATAAGAGGCTGACCGAGGG-3’, reverse primer 5’-CCCTCGGTCAGCCTCTTATGGAACAACGC-3’, HOIP C885A: forward Primer 5’-GCCCGAGGAGGCGCCATGCACTTTCACTGTACC-3’, reverse Primer 5’-GGTACAGTGAAAGTGCATGGCGCCTCCTCGGG-3’.

### Antibodies and reagents

The following antibodies were purchased and used according to the manufacturer’s instructions: anti-Myc (9E10) antibody (Covance, MMS-150P), anti-HA (HA.11 clone 16B12) antibody (Covance, MMS-101P), anti-Flag (M2) antibody (Sigma, F3165), anti-vinculin antibody (Sigma-Aldrich, V9131), anti-alpha-tubulin antibody (Abcam, ab15246), anti-ubiquitin (P4D1) antibody (Santa Cruz Biotechnology, sc-8017), anti-linear ubiquitin (LUB9) antibody (Life Sensors, #AB130), and anti-linear ubiquitin (LUB4) antibody (a kind gift from Japan Tobacco Inc. Pharmaceutical Frontier laboratories), anti-HOIP antibodies used for the detection of human HOIP (Aviva systems biology, ARP43241_P050, and Sigma, SAB2102031), anti-HOIL-1L antibody (Merck Millipore, MABC576), anti-SHARPIN antibody (Novus, NBP2-04116), anti-Fam105b/OTULIN antibody (Abcam, ab151117), anti-NEMO/IKKγ antibody (FL-419) (Santa Cruz, sc-8330), anti-IκB-α antibody (Cell Signaling, #4812), anti-pIκB-α antibody (Cell Signaling, #9246), anti-PARP antibody (Cell Signaling, #9542), anti-cleaved Caspase 3 antibody (Cell Signaling, #9664), anti-FADD antibody used for immunoprecipitation (Santa Cruz, sc-271748), anti-FADD antibody used for detection of FADD by immunoblotting (Abcam, ab124812), and anti-RIPK1 antibody (Cell signaling, #3493), and anti-phospho-RIPK1 antibody (Cell Signaling, #31122). A polyclonal antibody against mouse HOIP was raised against a recombinant protein containing a mouse HOIP fragment (aa 475-625) by immunizing rabbit (immunoGlobe, Germany). Secondary antibodies used for the immunoblotting are Goat anti-Mouse IgG-HRP (Bio-Rad, 170-6516) and goat anti-Rabbit IgG-HRP (Dako, P0448). Secondary antibodies used for immunoprecipitation were Protein G Agarose beads (Roche, 1124323301) and anti-FLAG (M2) beads (Sigma Aldrich, A2220)

Recombinant human Flag-TNF (Enzo, ALX-522-008-C050), recombinant human TNF (Peprotech, 300-01A), recombinant human TRAIL (Peprotech, 310-04), Lipopolysaccharide (LPS) (Sigma Aldrich, L4391), Cycloheximide (CHX) (Sigma Aldrich, C4859) and Z-Val-Ala-DL-Asp(Ome)-fluoromethylketone (z-VAD-fmk) (Bachem, N-1560) were also used. Recombinant Fc-Fas ligand was a kind gift from Pascal Schneider (Schneider et al., 1997).

### Tissue culture and transfection

Human embryonic kidney 293T (HEK293T) (ATCC) and immortalized MEFs were maintained at 37°C in 5% CO_2_ in Dulbecco’s modified Eagle’s medium (DMEM) (Sigma, D5648) supplemented with 10% fetal calf serum (ThermoFisher Scientific, 10270106), 1% L-glutamine (ThermoFisher Scientific, 25030-024), and 1% penicillin–streptomycin (Sigma, P0781). PCR-based mycoplasma tests confirmed all cells to be negative for mycoplasma contamination. Transfections in HEK293T or MEFs were performed using GeneJuice (Merck Millipore, 70967) according to the manufacturer’s protocol.

### Isolation and immortalization of mouse embryonic fibroblasts (MEFs)

Primary MEFs were isolated from E13.5 embryos (C57BL/6J *Hoip^+/+^* and *Hoip^K778R/K778R^*, C57BL/6J/KaLawRij *Hoip^+/+^; Sharpin^+/+^*, *Hoip^K778R/K778R^; Sharpin^+/+^*, *Hoip^+/+^; Sharpin^cpdm/cpdm^*, and *Hoip^K778R/+^; Sharpin^cpdm/cpdm^*) according to a standard protocol (Ikeda et al., 2011).

### Isolation of primary mouse dermal adult fibroblasts (MDFs)

Primary mouse dermal adult fibroblasts (MDFs) were isolated from ear tissue derived from mice at four weeks of age. Ear tissue was washed in 70% ethanol, air dried and minced into small pieces using a scalpel. Tissue pieces were collected in Dulbecco’s modified Eagle’s medium (DMEM) (Sigma, D5648) supplemented with 10% fetal calf serum (ThermoFisher Scientific, 10270106,) 1% penicillin–streptomycin (Sigma, P0781), Gentamicin (50µg/ml) (Thermofisher Scientific, 15750060), and 1% MEM non-essential amino acid solution (Thermofisher Scientific, 11140050) and centrifuged at 1,500 rpm for 5 minutes. Trypsin (Thermofisher scientific, 25300054) treated tissue pieces were incubated for 1 hour at 37°C, with a vortex step performed every 15 minutes. Fresh media was added and the tissue pieces were centrifuged, resuspended in fresh media and plated in a tissue culture dish. MDFs were seeded for cellular assays upon reaching confluency.

### Cell lysis

A method is described elsewhere (Ikeda et al., 2011). Briefly, cells were lysed in chilled lysis buffer (50mM HEPES (pH7.4) (Sigma Aldrich, H4034), 150mM NaCl, 1mM EDTA, 1mM EGTA, 1% Triton X-100, 10% Glycerol, 25mM NAF and 10µM ZnCl2, 1 x cOmplete protease inhibitor cocktail (Roche, 11836170001), 1mM PMSF (Roche, 10837091001) and 10mM NEM (Sigma-Aldrich, E3876) on ice. Lysates were cleared by centrifugation at 15,000 rpm for 15 minutes. For denaturing conditions, cells were lysed in 1% SDS-PBS and boiled at 96°C for 10 minutes as described before (Sasaki et al., 2015). Subsequently, lysates were sheared through a 27 3/4G needle (Becton Dickinson, BD 302200) several times, centrifuged at 15,000 rpm for 5 minutes (at room temperature) and the supernatant was subjected for further analysis.

### Immunoprecipitation

For immunoprecipitation of Myc-HOIP, anti-Myc antibody (1μg) was incubated for 2 hours at 4°C, followed by incubation with Protein G Agarose beads (Roche, 1124323301) (15μl) for 2 hours at 4°C. Beads were washed four times in lysis buffer. Proteins were eluted from beads using 30µl of 2X SDS sample buffer and heated at 96°C for 5 minutes.

A method for immunoprecipitation of the TNFR complex I is described in previous studies (Draber et al., 2015, Haas et al., 2009). Briefly, after serum starvation in 0.2% FCS-DMEM for 15 hours, MEFs (5-20×10^6^ cells) were treated with 1µg/ml of Flag-human TNF, washed by PBS twice, and lysed in 1ml of chilled IP-Lysis buffer (30mM Tris-HCI (pH7.4), 120mM NaCl, 2mM EDTA, 2mM KCI, 10% glycerol, 1%Trition X-100, 50mM NaF, 1 x cOmplete protease inhibitor cocktail (Roche, 11836170001), 1mM PMSF (Roche, 10837091001), 10mM NEM (Sigma-Aldrich, E3876), and 5mM NAVO_4_ (Sigma-Aldrich, S6508)) for 30 mins on ice. Lysates were centrifuged at 15,000 rpm for 30 minutes at 4°C. Flag-human TNF (1µg) was added to the 0hr control samples. After preclearing with Protein G Agarose beads for 1 hour at 4°C, anti-FLAG (M2) beads (Sigma Aldrich, A2220) (10µl of beads slurry) were incubated for 16 hours at 4°C, washed five times with the IP-Lysis Buffer and denatured for five minutes at 96°C in 2X SDS sample buffer.

A method for immunoprecipitation of TNFR complex/complex II is described elsewhere (Ang & Ting, 2018). Briefly, MEFs (5-20×10^6^) were treated with human TNF (100ng/ml), z-VAD-fmk (25µM) and cycloheximide (1µg/ml) for the indicated times and lysed in chilled DISC-IP buffer (150mM NaCl, 20mM Tris-HCL pH 7.5, 1mM EDTA, 0.2% NP40, 10% glycerol supplemented with cOmplete protease inhibitor cocktail (Roche, 11836170001), 0.1mM Na_3_VO_4_ (Sigma-Aldrich, S6508), 100mM NEM (Sigma-Aldrich, E3876), 1mg/ml of BSA (VWR International, 422351S) for 10 minutes on ice. Lysates were centrifuged at 15,000 rpm for 15 mins and supernatant was precleared with Protein G Agarose beads (25µl) for 1.5 hours at 4°C, followed by immunoprecipitation with α-FADD antibody (2µg) (Santa Cruz; sc-271748) incubation for 16 hours at 4°C. Subsequently, Protein G Agarose beads (25µl) incubation for 1.5 hours at 4°C. Immunoprecipitated samples were washed four times with DISC-IP buffer. Samples were heated at 70°C for 20 minutes in 2X SDS sample buffer.

### GST-Linear TUBE pulldown

GST-Linear TUBE pulldown was performed as previously described (Asaoka et al., 2016). Briefly, cells were lysed in mammalian lysis buffer on ice and cleared by centrifugation. GST-empty and GST-Linear TUBE immobilized on glutathione Sepharose 4B beads were incubated with supernatants for 12 hours at 4°C. Pulldown samples were washed five times in lysis buffer, and heated at 96°C for 5 minutes in 2X SDS sample buffer.

### Immunoblotting

The protocol used for immunoblotting was described previously (Ikeda et al., 2011) (Kumari et al., 2014). Briefly, samples were resolved by SDS-PAGE, and transferred to a nitrocellulose membrane (GE Healthcare, 10600019 or 10600001). The membrane was proceeded with Ponceau S (Roth, 5938.1) staining to monitor the transferred proteins. Membranes were washed, blocked with 5% BSA-TBS, and blotted with the indicated primary antibodies diluted in 5% BSA-TBS at 4°C overnight. Subsequently, membranes were incubated with a secondary antibody according to the manufacturer’s instructions, and signal was detected with Western Blotting Luminol Reagent (Santa Cruz; sc-2048) on high-performance chemiluminescence films (GE Healthcare, Amersham Hyperfilm ECL, 28906837).

### Luciferase-based NF-κB gene reporter assay

HEK293T cells were seeded in 96-well plates (1×10^4^ cells/well) (Thermoscientific, 136101) and transfected with pNF-κB-Luc (Stratagene) and phRK-TK (Renilla) (Promega) using GeneJuice (Merck Millipore, 70967). After 48 hours, samples were subjected to a luciferase assay using the Dual-Glo Luciferase Assay System (Promega; E2940) according to the manufacture’s protocol. Luciferase and Renilla signal were measured by the Synergy H1 hybrid multimode microplate reader (BioTek) and monitored by Gen5 software. Each experimental sample was carried out in quadruplicate and normalized to the Renilla signal.

### Protein Purification

A method is described elsewhere (Asaoka et al, 2016; Ikeda et al, 2011). Briefly, plasmids were transformed into BL21 (DE3) *E.coli.* Bacterial cells were grown in the LB media at 37°C. Expression of GST tagged fusion proteins were induced using 100μM IPTG (Thermoscientific, R0392) at OD_600_= 0.8 in 4-6 litres of culture. 100µM ZnCl_2_ (Sigma-Aldrich, 229997) was added during induction of HOIP, HOIL-1L and SHARPIN expression only. Cultures were grown overnight at 18°C. Cells were centrifuged and resuspended in the suspension buffer (100mM HEPES (Sigma Aldrich, H4034), 500mM NaCl, 1mM TCEP-HCl (ThermoScientific, 20491) pH 7.4 which was supplemented with recombinant DNase I (1000U) (Roche,04536282001), cOmplete protease EDTA-free inhibitor cocktail (Roche, Roche, 11836170001) and 1mM PMSF (100mM in isopropanol, Roche, 10837091001). Cells were sonicated and 0.5% Triton X-100 was added to the lysate. The lysate was cleared by centrifugation and applied to a 5ml GSTrap FF column (GE Healthcare, 17513101) to initially purify GST-proteins. The GST-tag was removed by overnight on-column cleavage with the PreScission Protease (homemade). Protein eluates were further resolved using size exclusion chromatography on gel filtration columns using the Superdex 200 (16/600) (GE Healthcare, GE28-9893-35) or Superdex 75 (16/600) (GE Healthcare, GE28-9893-33) in a buffer containing 50mM HEPES (Sigma Aldrich, H4034), 150mM NaCl, 1mM TCEP-HCl (ThermoScientific, 20491), pH 7.4. Eluted fractions were analysed in SDS-PAGE stained with InstantBlue^TM^ Protein Stain (Expedeon, 1SB1L) and the fractions containing the desired protein were pooled together. Proteins were concentrated using a Vivaspin concentrator (Sartorius) with a half lower MWCO than the size of the protein being purified. Protein concentrations were determined by UV absorption at 280 nm using calculated extinction coefficients or compared to known BSA standards visualized by SDS-PAGE and stained with Instant Blue. A baculovirus for insect expression of His_6_-mouse Ube1 in Hi5 cells was a kind gift from Kazuhiro Iwai and was expressed and purified as previously described (Iwai, Yamanaka et al., 1999).

### NanoLC-MS Analysis

Samples containing HOIP were separated using SDS-PAGE using 4-15% Mini-PROTEAN TGX gels (Bio-Rad; #4561083). Gels were silver stained according to Blum’s protocol (Helmut et al., 1987). Gel fragments containing the HOIP band and above were extracted from the gel. Following this, the gel bands were reduced, alkylated and digested with Trypsin.

The nano HPLC system used was an UltiMate 3000 RSLC nano system (Thermo Fisher Scientific, Amsterdam, Netherlands) coupled to a Q Exactive Plus mass spectrometer (Thermo Fisher Scientific, Bremen, Germany), equipped with a Proxeon nanospray source (Thermo Fisher Scientific, Odense, Denmark). Peptides were loaded onto a trap column (Thermo Fisher Scientific, Amsterdam, Netherlands, PepMap C18, 5 mm × 300 μm ID, 5 μm particles, 100 Å pore size) at a flow rate of 25 μL min-1 using 0.1% TFA as mobile phase. After 10 min, the trap column was switched in line with the analytical column (Thermo Fisher Scientific, Amsterdam, Netherlands, PepMap C18, 500 mm × 75 μm ID, 3 μm, 100 Å). Peptides were eluted using a flow rate of 230 nl min-1 and a binary 1h gradient, respectively 105 min.

The gradient starts with the mobile phases: 98% A (water/formic acid, 99.9/0.1, v/v) and 2% B (water/acetonitrile/formic acid, 19.92/80/0.08, v/v/v), increases to 35%B over the next 60 min, followed by a gradient in 5 min to 90%B, stays there for 5 min and decreases in 5min back to the gradient 98%A and 2%B for equilibration at 30°C.

The Q Exactive Plus mass spectrometer was operated in data-dependent mode, using a full scan (m/z range 380-1650, nominal resolution of 70,000, target value 3E6) followed by MS/MS scans of the 12 most abundant ions. MS/MS spectra were acquired using normalized collision energy of 27%, isolation width of 2.0 m/z, resolution of 17.500 and the target value was set to 1E5. Precursor ions selected for fragmentation (exclude charge state 1) were put on a dynamic exclusion list for 10 s. Additionally, the intensity threshold was calculated to be 4.0E4. The peptide match feature and the exclude isotopes feature were enabled.

### Data Processing protocol for analyzed peptides

For peptide identification, the RAW-files were loaded into Proteome Discoverer (version 1.4.1.14, Thermo Scientific). All hereby created MS/MS spectra were searched using MSAmanda v1.4.14.7870 (Dorfer V. et al., J. Proteome Res. 2014 Aug 1;13(8):3679-84). The RAW-files were searched against the Swissprot sequence database, using the taxonomy human (20,171 sequences; 11,317,551 residues). The following search parameters were used: Beta-methylthiolation on cysteine was set as a fixed modification, oxidation on methionine, deamidation on asparagine and glutamine, acetylation on lysine, phosphorylation on serine, threonine and tyrosine and ubiquitination on lysine were set as variable modifications. Monoisotopic masses were searched within unrestricted protein masses for tryptic enzymatic specificity. The peptide mass tolerance was set to ±5 ppm and the fragment mass tolerance to 0.03 Da. The maximal number of missed cleavages was set to 2. The result was filtered to 1 % FDR on protein level using Percolator algorithm integrated in Thermo Proteome Discoverer. The localization of the post-translational modification sites within the peptides was performed with the tool ptmRS, based on the tool phosphoRS (Taus T. et al., J. Proteome Res. 2011, 10, 5354-62). Peptide areas have been quantified using in-house-developed tool APQuant (publication under review).

### *In vitro* ubiquitination assays

A method for in vitro ubiquitination assay is described elsewhere (Asaoka et al., 2016). Briefly, recombinant proteins of Ubiquitin (10µg) (Sigma-Aldrich, U6253) or human His_6_-ubiquitin (Boston Biochem, U-530), mouse E1 (Ube1) (150ng), human UbcH7 (300ng), human HOIP (5µg) or human HOIP UBA-RBR-C (aa 476-1072) (5µg), human SHARPIN (1µg), human HOIL-1L (1µg), human NEMO (5µg) and ATP (2mM) (Roche, 1051997900) were incubated in a reaction buffer consisting of 50mM HEPES (Sigma Aldrich, H4034) (pH7.5), 150mM NaCl, 20mM MgCl_2_ for the indicated times at 37°C. Reactions were terminated by 2x SDS sample buffer at 96°C for 1 minute. Samples were subjected to SDS-PAGE and subsequent immunoblotting.

### *In vitro* deubiquitination assays (UbiCRest assays)

Deubiquitination assays were performed as previously described (Hospenthal et al., 2015). Briefly, recombinant deubiquitinases (3µM for vOTU (CCHFV OTU domain, aa1-183), 10µM for human OTULIN, 3µM for human USP21 (USP domain, aa 196-565) were activated in activation buffer (150mM NaCl, 25mM Tris pH7.5, and 10mM DTT) for 10 minutes at room temperature. Subsequently, samples in 10x DUB reaction buffer (500mM NaCl, 500mM Tris pH7.5, and 50mM DTT) were incubated with activated DUB for 30 minutes at 37°C. Reaction was terminated by 2X SDS sample buffer.

### Bioinformatics analysis

A multiple sequence alignment of HOIP orthologues was performed with MAFFT (Katoh & Toh, 2008) (Asaoka et al., 2016) and visualized using Jalview (Waterhouse et al., 2009).

### qRT-PCR

A method is described elsewhere (Asaoka et al., 2016). 3 x10^5^ MEFs or 1×10^6^ primary BMDMs in 6-well plates were serum starved for 15 hours in 0.2% FBS-DMEM, treated with human TNF (20ng/ml) for the indicated timepoints. Samples washed by chilled PBS two times and the total RNA was extracted using TRIzol (Life Technologies, 15596018), treated with the TURBO DNA-free kit (Invitrogen, AM1907). 500ng of RNA from MEFs or 350ng of RNA from BMDMs were reverse transcribed using oligo(dT)18 primer (New England Biolabs, #513165) and SuperScript II Reverse Transcriptase (Invitrogen, 18064-014). The cDNA was proceeded by using a standard qPCR method with GoTaq qPCR master mix (Promega, A6002) and the CFX96 BioRad CFX 96 Real-Time PCR detection system. β-actin was used for normalization. Analysis was carried out using the 2^-ΔΔCt method. The sequences of primers used against mouse genes are following.

IκBα: forward primer 5’-GCTGAGGCACTTCTGAAAGCTG-3’, reverse primer 5’-TGGACTGGCAGACCTACCATTG-3’, ICAM: forward primer 5’-AAGGAGATCACATTCACGGTG-3’, reverse primer 5’-TTTGGGATGGTAGCTGGAAG-3’, VCAM: forward primer 5’-CTGGGAAGCTGGAACGAAGT-3’, reverse primer 5’-GCCAACACTTGACCGTGAC-3’, A20: forward primer 5’-AAAGGACTACAGCAGAGCCCAG-3’, reverse primer 5’-AGAGACATTTCCAGTCCGGTGG-3’, β-actin: forward primer 5’-CGGTTCCGATGCCCTGAGGCTCTT-3’, reverse primer 5’-CGTCACACTTCATGATGGAATTGA-3’.

### Caspase 8 assay

5×10^4^ cells/well in a 96-well white plate (Thermoscientific, 136101) were treated by human TNF (100ng/ml), with or without cycloheximide (1µg/ml) and z-VAD-fmk (20µM). Caspase 8 activity was measured by using the Caspase Glo 8 assay system (Promega, G8202) according to the manufacturer’s protocol.

### Generation of C57BL/6J *Hoip^K778R/K778R^* knockin mice

A method is described as previously (Wang et al., 2013). Briefly, the gRNA was designed using the online tool (crispr.mit.edu). Annealed oligonucleotide with gRNA targeting sequence was cloned into px330 plasmid by a standard subcloning method (Addgene plasmid, #42330, a gift from Feng Zhang (Cong et al., 2013)). The T7-gRNA product amplified by PCR was used as the template for *in vitro* transcription using the MEGAshortscript T7 kit (Invitrogen, AM1345). *In vitro* transcribed gRNA purified using the MEGAclear kit (Invitrogen, AM1908), Cas9 mRNA (Sigma CAS9MRNA-1EA), single strand oligonucleotide donor template containing K778R mutation, a silent mutation of the PAM sequence and a silent XmnI restriction site (ssOligo) (5’-TGCATCTTGTTCCCAGCTCAGAGAGAGCCTAGACCCCGATGCATATGCCCTGTTTCACAAGAGGCTGACCGAAGCTGTTCTTATGCGAGACCCCAAGTTCTTGTGGTGCGCCCAGGTAAACCTGACAAACAGAGTGAACT-3’) were used for injection. Superovulation-induced female C57BL/6J donor mice (3-5 weeks old), treated with 5IU of pregnant mare’s serum gonadotropin (PMSG) (Hölzel Diagnostika, OPPA01037) and subsequently with 5IU of human chorionic gonadotropin (hCG) (Intervet, GesmbH) were mated and zygotes were isolated in M2 media (Merck Millipore, MR-015P-D) and cultured in KSOM medium (Cosmo Bio Co., Ltd, R-B074). The microinjection mix(100ng/µl of Cas9 mRNA, 50ng/µl of gRNA, 200ng/µl of ssOligo) was microinjected into the cytosol of zygotes followed by transfer to pseudo-pregnant females. The genomic fragment of targeted region in *Hoip^K778R/K778R^* knockin founder mice was confirmed by Sanger sequencing.

### Genotyping of *Hoip^K778R/K778R^* knockin mice

A PCR amplified genomic DNA fragment using forward primer 5’-CGATCCTCTTGCCTCCATGT-3’ and reverse primer 5’-CCAGCTGTTCGCGTTCATA-3’ was digested with XmnI (NEB; R0194L). Undigested and digested samples were proceeded for electrophoresis using 2% agarose gels.

### Mouse husbandry

*Hoip^K778R/K778R^* knockin C57BL/6J mice, *Sharpin^cpdm/cpd^*^m^ C57BL/KaLawRij (Ikeda et al., 2011), and *Tnfrsf1a^tm1Mak^*/*TNFRp55-deficient* C57BL/6J mice (JAX stock #002818) were used in this study. All animal procedures were conducted in accordance with European, Austrian and institutional guidelines and protocols. All animal conduct was approved by local government authorities.

### Histopathological analysis

A method is described elsewhere (Kumari et al., 2014). Briefly, mouse tissues of dorsal and ventral skin, lung, spleen and liver, kidney, small intestine, cecum and colon were fixed in 10% neutral buffered formalin (Sigma, HT501128), processed with a microwave hybrid tissue processor (LOGOS, Milestone Medical), embedded in paraffin, sectioned (Microm, HM 355) for the hematoxylin and eosin staining in an automated stainer (Microm HMS 740). Immunohistochemistry was performed using an automated immunostainer (Bond III, Leica). Primary antibodies used for immunohistochemistry are KRT14 (Sigma Aldrich, SAB4501657, 1:200) CASP3, cleaved (Cell Signaling, 9661, 1:100) F4.80 (Bio Rad, MCA497G, 1:100), Ly6G (Abcam, ab2557, 1:500) and secondary antibodies used are rabbit anti-rat IgG (Abcam, ab6733, 1:500), goat anti-rabbit IgG (Dako, E0432 1:500). Signal was detected with the Leica Bond Intense R Detection system. Slides were evaluated by a board certified veterinary comparative pathologist with a Zeiss Axioskop 2 MOT microscope and images were acquired with a SPOT Insight color camera (SPOT Imaging, Diagnostic Instruments, Inc.).

For the skin thickness measurement, digital images of sections stained with an KRT14 antibody were taken by the 3D Histech *Pannoramic Flash III* whole slide scanner, evaluated with the *Pannoramic Viewer* software. From each digital scene, 10 non-contiguous, representative foci were selected from a region spanning 4 mm. In each focus, the thickness of the entire epidermis, keratin layer and squamous cell layer were measured. Measurements were tabulated and the average of ten measurements was recorded per sample.

### Mouse embryo analysis

Female mice mated with male mice in breeding age were checked for copulation plugs daily. The embryos were collected at different embryo stages (E10.5, E12.5, E13.5), and fixed in 10% neutral buffered formalin (Sigma, HT501128) for 12-24 hours at room temperature. Fixed embryos washed with PBS were imaged using a Lumar-Florescence Stereomicroscope with a color SPOT camera at 9X magnification. Embryos were processed, embedded, sectioned and stained with hematoxylin and eosin for histopathological analysis.

### Statistical Analysis

Analysis was performed by the GraphPad Prism 8 software (GraphPad Software, Inc) and the mean values with standard deviation are shown. A One-way ANOVA test and Tukey’s post hoc test were used. The significance and confidence level were set at 0.05 and P values are indicated in each figure legends.

## Acknowledgements

We thank Merle Hantsche-Grininger (EMBL, Heidelberg) and Michaela Morlock (2016 VBC summer student, currently AstraZeneca, Göteborg, Sweden) for their initial contribution to the project setups, Katrin Rittinger (Crick Institute, London) and Paul Elliott (University of Oxford, Oxford) for discussions on the structural aspect of HOIP, and all the Ikeda Lab members for constructive discussions and suggestions on the project, as well as the team work. We also thank Anna Szydłowska and Kikue Tachibana (IMBA, Vienna) for the technical advice on CRISPR-Cas9-based knockin mouse generation, Adrian Ting (Icahn School of Medicine at Mount Sinai, New York) for an advice on ear-derived fibroblast isolation, Richard Imre (Protein Chemistry, Vienna) for the Mass spectrometry data analysis, the IMP-IMBA core facilities of transgenic service, comparative medicine, molecular biology service, biooptics, and VBCF facilities, ProTech and Histo Pathology for their technical assistance. Research in Ikeda Lab is supported by ERC Consolidator Grant (LUbi, 614711), FWF stand-alone grant (P 2550 8) and Austrian Academy of Sciences. We also thank Angela Andersen from the Life Science Editors for editing the manuscript.

## Author contributions

L.M.F. preformed most of the experiments, analyzed data, made figures and contributed in writing the manuscript. L.D. expressed and purified recombinant proteins required for *in vitro* assays, A.S. analyzed amino acid sequences of HOIP orthologues and made alignment, K.M. contributed to the mass spectrometry analysis, and A.K. performed histological analysis. F.I. planned and conducted the project, analyzed data, made figures, and wrote the manuscript.

## Supplementary figure legends

**Supplementary Figure 1.**
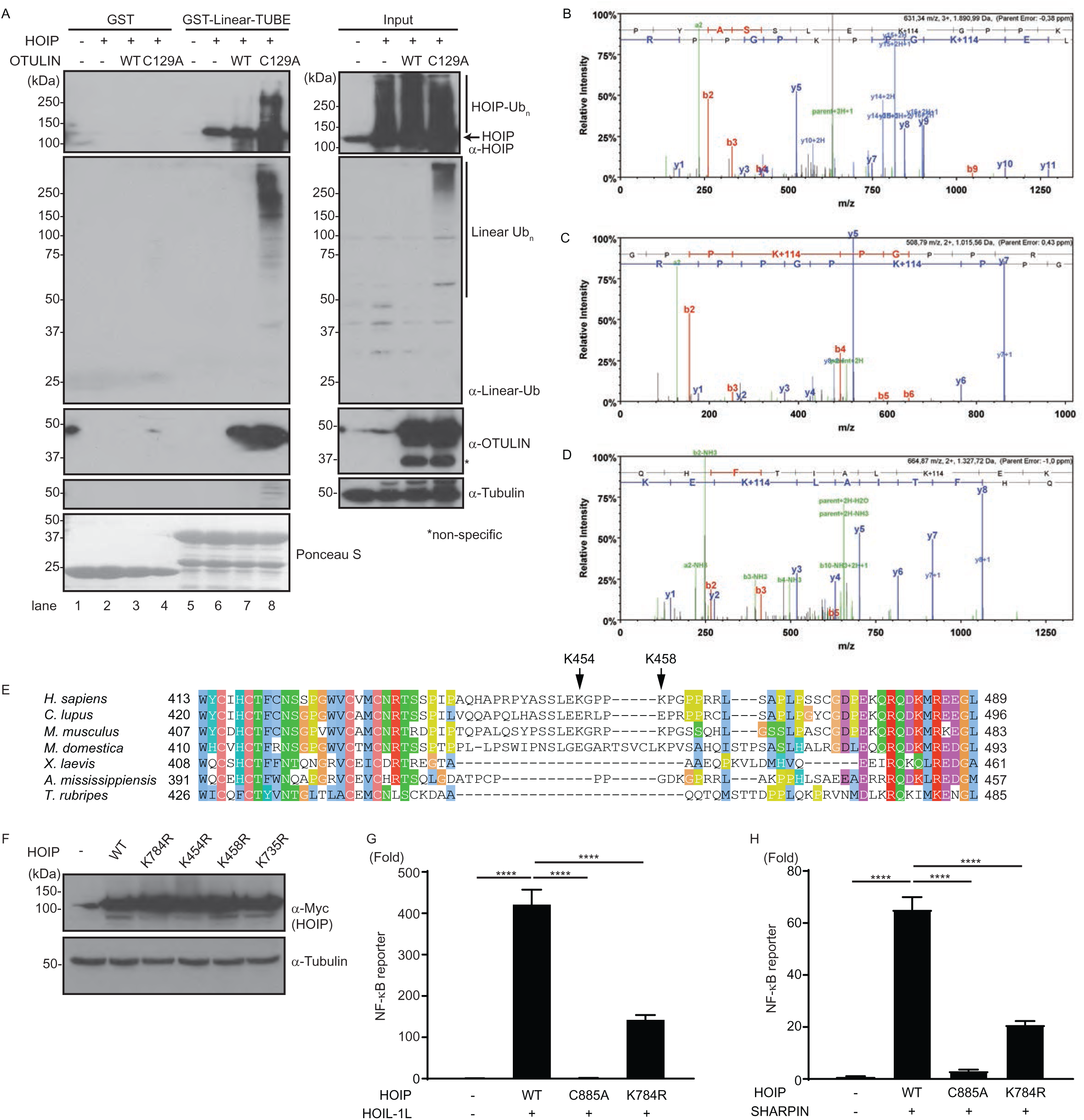
HOIP is ubiquitinated in cells. A. Immunoblotting of ubiquitinated HOIP enriched by Linear-TUBE pulldown assays in total cell extracts from HEK293T cells transiently expressing Myc-HOIP, Myc-OTULIN wildtype (WT) or a catalytically inactive mutant (C129A). Linear ubiquitin chains and ubiquitination of HOIP were detected by immunoblotting. Representative data shown from three independent experiments. B-D. Mass spectrometry spectra corresponding to a peptide containing HOIP-K454, K458 and K735 with double Gly (114+K). E. Multiple sequence alignment of HOIP orthologues of the indicated species illustrating the position of K454 and K458 in HOIP according to the ClustalX color scheme. F. Expression of Myc-HOIP mutants of the identified ubiquitination sites (K784, K454, K458, K735) examined by immunoblotting. Total cell extracts of HEK293T cells transiently expressing Myc-HOIP WT or mutants subjected to SDS-PAGE followed by immunoblotting using the indicated antibodies. An anti-Tubulin antibody used for loading control. G, H. NF-κB activation examined by reporter gene assays in HEK293T cells. Myc-HOIP WT, Myc-HOIP C885A or Myc-HOIP K784R were expressed in HEK293T cells with HOIL-1L-HA or FLAG-SHARPIN with reporter plasmids. Luciferase signal was normalized by internal control, Renilla signals. Representative data shown from three independent experiments, n=4, data presented as mean ± SD, ****p≤0.001 by ANOVA.

**Supplementary Figure 2.**
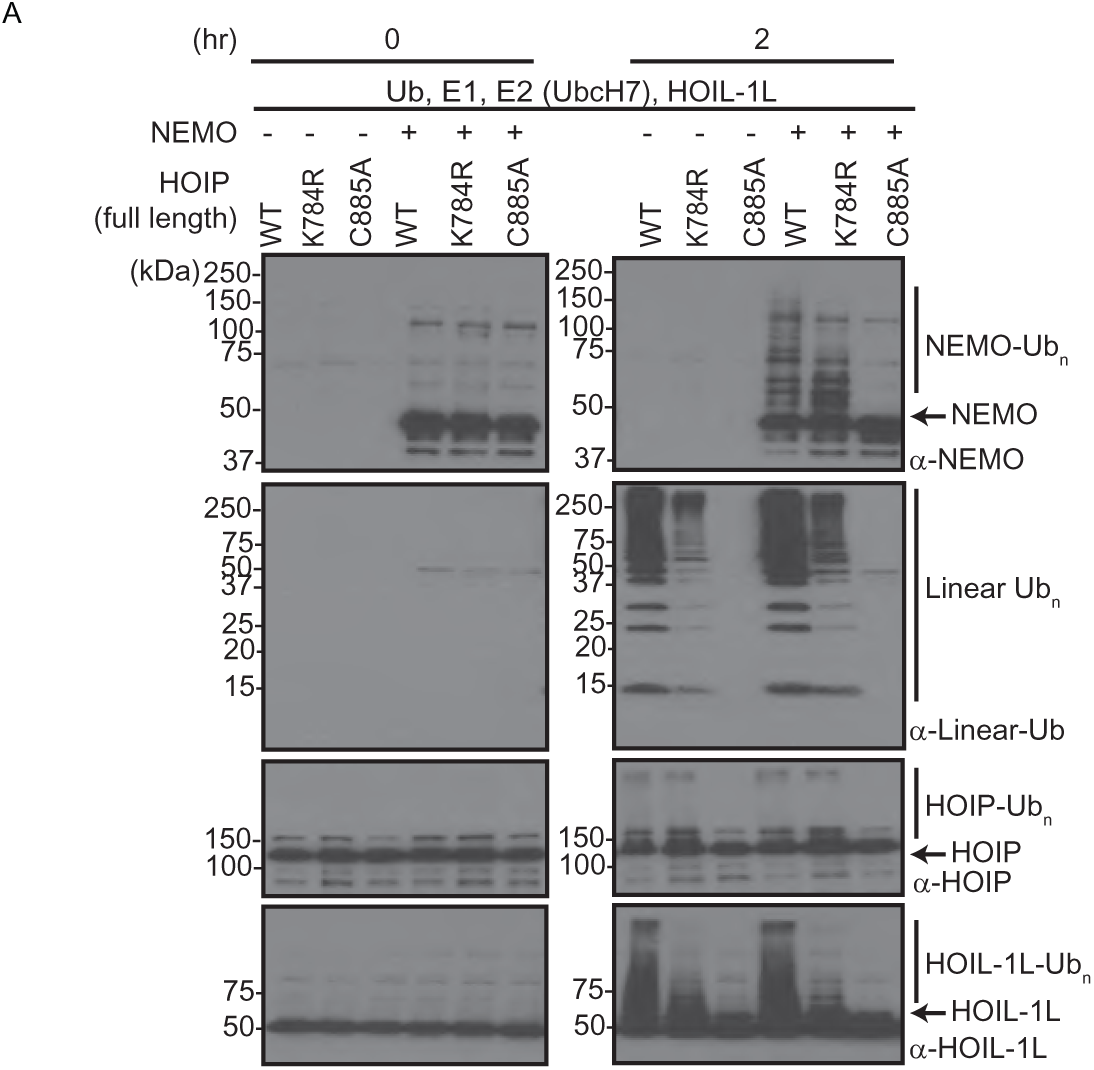
Recombinant HOIP K784R protein catalyzes linear ubiquitin chains *in vitro*. A. Immunoblotting of i*n vitro* ubiquitination assay samples using recombinant proteins of ubiquitin, E1, E2 (UbcH7), E3 (LUBAC components, HOIP and HOIL-1L without SHARPIN) and a known substrate NEMO. HOIP wildtype (WT), K784R or C885A mutants were used as indicated. Unanchored linear ubiquitin chain formation, ubiquitination of NEMO, HOIP and HOIL-1L were detected by immunoblotting using the indicated antibodies. Representative data shown from three independent experiments.

**Supplementary Figure 3.**
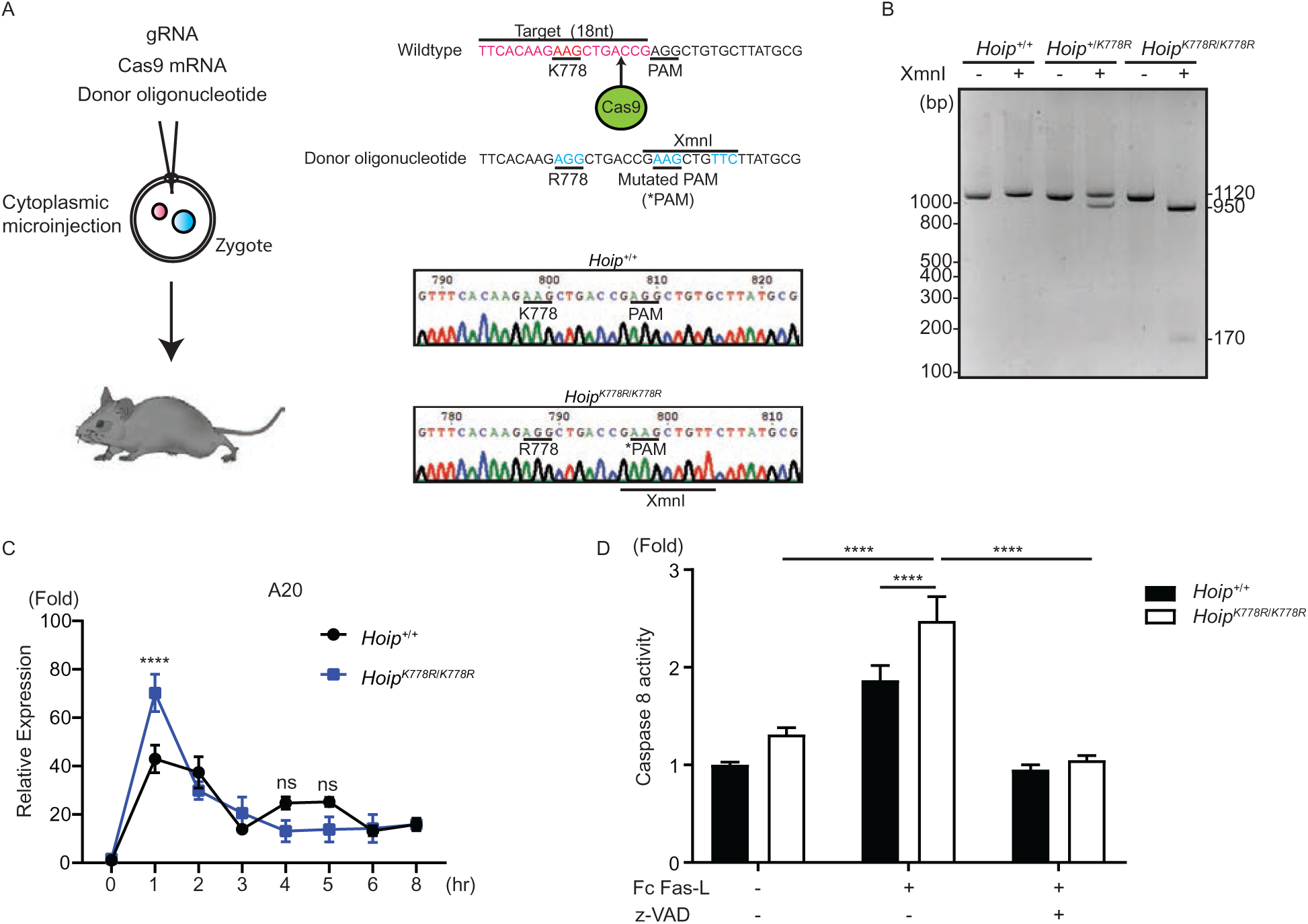
Generation of *Hoip^K778R/K7784R^* knockin mice by CRISPR/Cas9 and cellular responses. A. A schematic illustrating the strategy to generate *Hoip^K778R/K7784R^* knockin mice. Guide RNA (gRNA), Cas9 mRNA, and donor single-stranded oligonucleotide were microinjected into the cytosol of zygotes. Mouse HOIP K778 (equivalent to human HOIP K784) was mutated to Arg (R778), a silent mutation for an Xmn1 restriction site was introduced for genotyping, as well as for the PAM sequence. Sanger sequencing results of the corresponding target region are shown. B. Genotyping of *Hoip^+/+^*, *Hoip^+/K778R^* and *Hoip^K778R/K778R^* mice. Restriction Fragment Length Polymorphism (RFLP) analysis of a PCR product (1120bp) of genomic DNA isolated from *Hoip^K778R^* offspring using Xmn1. *Hoip^+/+^*: undigested, *Hoip^+/K778R^*: one allele digested, *Hoip^K778R/K778R^*: both allege digested. C. Induction of TNF-dependent NF-κB target genes, A20 in immortalized MEFs (*Hoip^+/+^* and *Hoip^K778R/K778R^*) examined by qRT-PCR. RNA extracted from MEFs treated with hTNF (20ng/ml) for the indicated time followed by RT, subjected to qPCR. Signals were normalized to β-actin. Representative data shown from three independent experiments (n=3). D. Caspase 8 activity in FasL-treated *Hoip^+/+^* and *Hoip^K778R/K778R^* MEFs. A luminol-based Caspase 8 assay data using MEFs treated by Fc-FasL (200ng/ml) for two hours. A representative data shown from two independent experiment, n=4. Data presentation: mean ± SD, ****p≤0.001 by ANOVA.

**Supplementary Figure 4.**
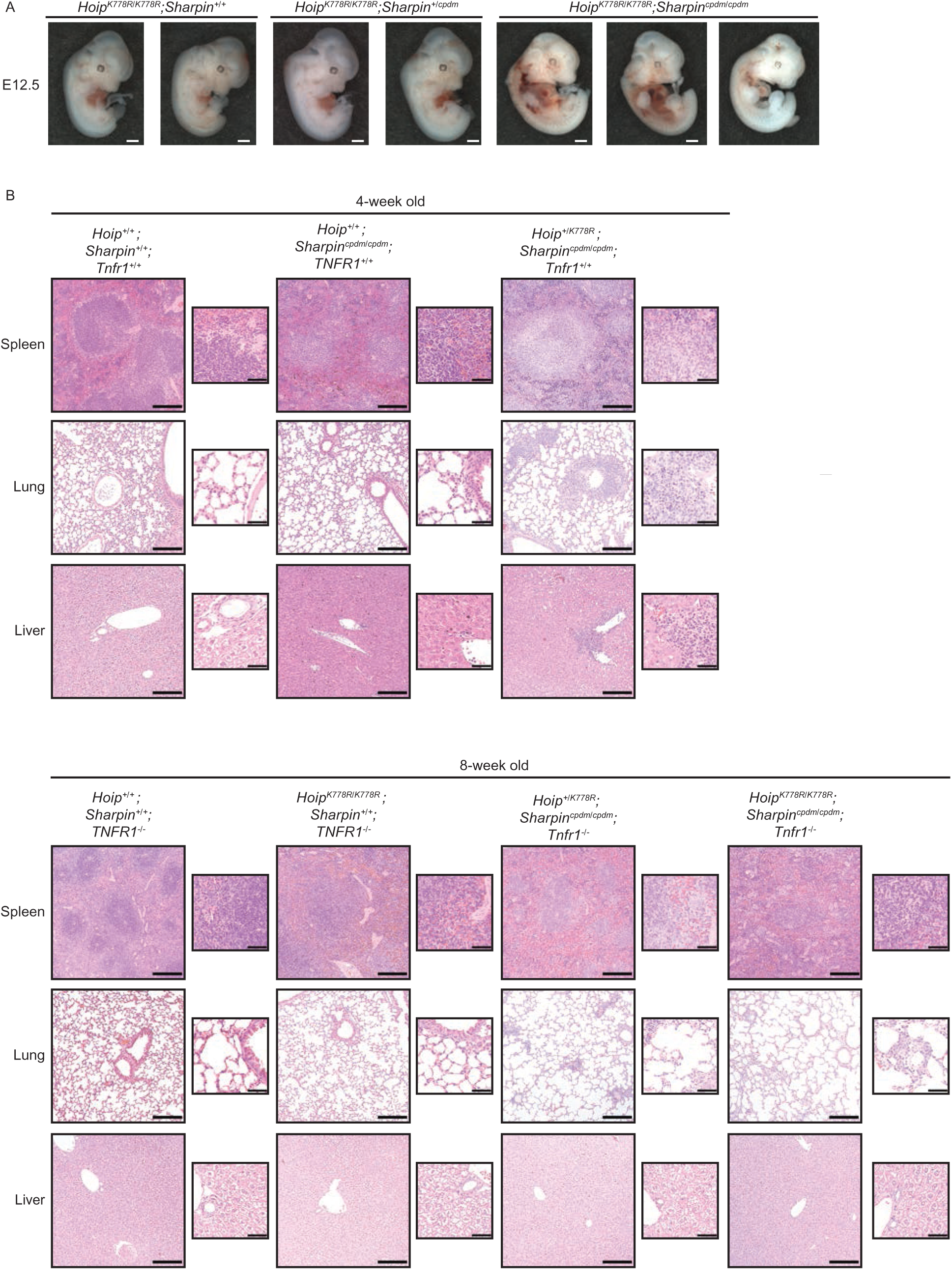
Grossly abnormal *Hoip^K778R/K778R^;Sharpin^cpdm/cpdm^* embryos and mitigation of systemic inflammation in *Hoip^K778R/K778R^;Sharpin^cpdm/cpdm^* by TNFR1 knockout. A. Gross images of *Hoip^K778R/K778R^;Sharpin^+/+^*, *Hoip^K778R/K778R^;Sharpin^+/cpdm^* and *Hoip^K778R/K778R^;Sharpin^cpdm/cpdm^* embryos at E12.5. Scale bars:1mm B. H&E staining of spleen, lung and liver sections from mice of the indicated genotypes. Scale bars: 200μm and 50μm (higher magnification images).

**Supplementary Figure 5.**
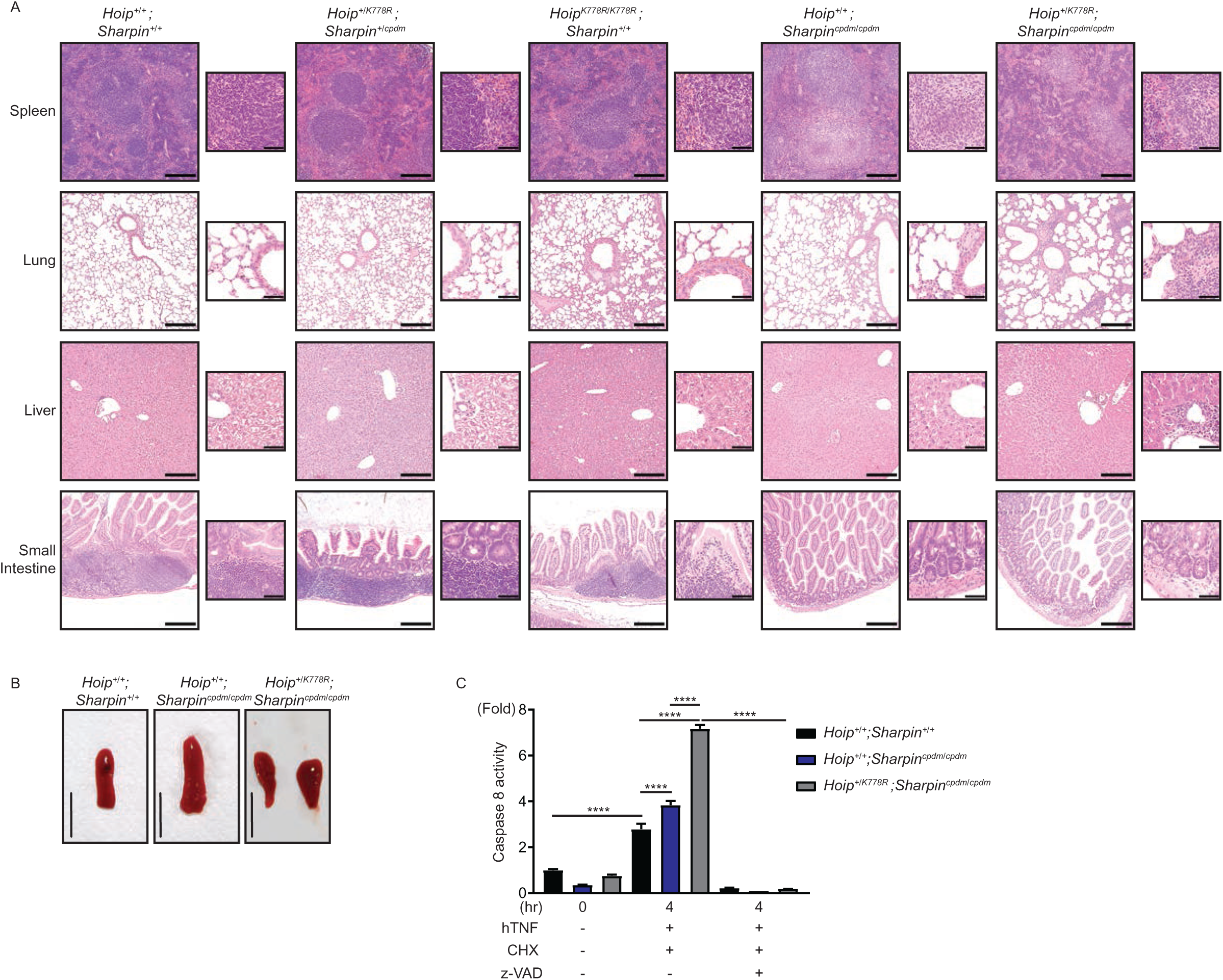
*Hoip^+/K778R^; Sharpin^cpdm/cpdm^* mice show multi-organ inflammation. A. H&E staining of spleen, lung, liver and small intestine from mice of the indicated genotypes at four weeks of age. Scale bars: 200μm and 50μm (higher magnification images). B. Macroscopic images of spleens from mice of the indicated genotypes at four weeks of age. Two representative spleens are shown for *Hoip^+/K778R^; Sharpin^cpdm/cpdm^* mice. Scale bars: 10mm. C. TNF-induced Caspase 8 activity in primary ear-derived fibroblasts from mice of the indicated genotypes. Luminol-dependent activity of Caspase 8 in MEFs treated with hTNF (100ng/ml) with or without CHX (1µg/ml) or z-VAD (20µM). Representative data from three independent experiments (n=4, data presented as mean ± SD, ****p≤0.0001).

